# Impacts of nucleosome positioning elements and pre-assembled chromatin states on expression and retention of transgenes

**DOI:** 10.1101/2024.08.21.607776

**Authors:** Ronard Kwizera, Junkai Xie, Nathan Nurse, Chongli Yuan, Ann L Kirchmaier

## Abstract

Transgene applications ranging from gene therapy to development of stable cell lines and organisms rely on maintaining expression of transgenes. To-date, the use of plasmid-based transgenes has been limited by loss of their expression shortly after delivery into target cells. This short-lived expression of plasmid-based transgenes has been largely attributed to host cell-mediated degradation and/or silencing of transgenes. To assess the impact of “priming” plasmid-based transgenes to adopt accessible chromatin states to promote gene expression, nucleosome positioning elements were introduced at promoters of transgenes, or vectors were pre-assembled into nucleosomes containing unmodified histones or histone mutants mimicking constitutively acetylated states at residues 9 and 14 of histone H3 or residue 16 of histone H4 prior to their introduction into cells, then transgene expression was monitored over time. DNA sequences capable of positioning nucleosomes could positively impact expression of adjacent transgenes in a distance-dependent manner in the absence of pre-assembly into chromatin. Intriguingly, pre-assembly of plasmids into chromatin facilitated prolonged expression of transgenes relative to plasmids that were not pre-packaged into chromatin. Interactions between pre-assembled chromatin states and nucleosome positioning-derived effects on expression of reporter genes were also assessed and, generally, nucleosome positioning played the predominant role in influencing gene expression relative to priming with hyperacetylated chromatin states.

## 1. Introduction

Effective expression of transgenes for a variety of applications ranging from generation of cell lines that stably express a protein of interest for basic research or industrial purposes to use in gene therapy relies on long-term expression of transgenes at appropriate levels. Currently, gene therapy strategies are predominantly applied through engineered integrating or non-integrating viral vehicles that are based on adenovirus, adeno-associated virus, or retroviruses [1-5]. However, several challenges accompany viral-based gene therapy, including immunogenicity, and cost [4,6,7]. Moreover, when transgene vectors become integrated into the host cell genome, their transgenes may be subject to position effect variegation where chromatin states adjacent to sites of integration are adopted, often resulting in the transgene becoming silenced [8-10]. Such issues motivate the development of alternate approaches, including application of nonviral plasmid-based vectors. However, one major challenge limiting effective use of nonviral plasmids has been rapid loss of expression of the encoded transgene following delivery into cells and organisms [11-13]. This can occur through rapid loss of the plasmid DNA itself from proliferating cell populations in the absence of selection [14-17], or through the adoption of chromatin states incompatible with efficient gene expression [18-20]. Like the host cell genome, exogenous episomal DNAs delivered into cells undergo host cell-mediated packaging into chromatin and acquire post-translational modifications in that chromatin that can either promote or silence expression of transgenes over time [21,22].

Transcription of DNA within chromatin is influenced by post-translational modifications on histones by controlling accessibility to DNA promoter regions via relaxation or compaction of chromatin or via recruitment of transcriptional machinery [23-27]. For example, acetylation of lysine residues on H3 and H4 neutralizes their positive charges, thus reducing their affinity for the negatively charged DNA, thereby relaxing nucleosomes in acetylated chromatin [28,29]. Histone acetylation correlates with active gene expression, whereas deacetylation, which restores positive charges to lysine residues, correlates with low levels of gene expression or gene silencing [30,31]. Consistent with this trend, acetylated H3K9 and H3K14 are enriched at promoters of actively transcribed genes [32,33]. Also, acetylated of histone H4K16 promotes the formation of relaxed chromatin via disrupting inter-nucleosome interactions and thus, can enhance accessibility of DNA within chromatin [29]. In contrast to acetylation, other modifications promote silencing of gene expression. For example, methylation of H3K9 by SUV39H1 both prevents H3K9 acetylation and promotes gene silencing via serving as a binding site for HP1, a structural component of heterochromatin [24,34-36].

Successful long-term application of nonviral, plasmid-based transgene vectors will require the establishment and maintenance of chromatin states that promote access of the transcriptional machinery to regulatory elements at genes and prolonged expression of those transgenes. Several strategies aimed at overcoming challenges associated with delivery and expression of transgenes have been explored previously, including the application of enhancer elements to facilitate expression [37-39], or of insulator elements (e.g. ubiquitous chromatin opening elements, UCOEs, and stabilizing anti-repressor elements, STAR) to protect transgenes from host cell-mediated silencing [40-42]. While insulator-based strategies can improve transgene expression, they have also been associated with a decline in expression following prolonged culturing and dramatic decreases in vector copy numbers shortly after delivery [43,44], possibly due to increased transgene expression causing toxicity [45-47]. Nuclear delivery of expression vectors can be enhanced via polyplexes with peptides based on the N-terminal tail of histone H3, which interacts with the nuclear import receptor Importin 4 [48-50]. Use of exogenous individual histone-plasmid DNA complexes have also been explored as methods of gene transfer, but have been limited in their success (for H3, see [51], see also [52]). However, in these systems, the charge-charge-based complexes formed between individual histones or histone fragments and DNA do not reflect natural packaging of DNA into nucleosomes.

Here, we assessed how transgene expression from the human elongation factor 1a (EF1α) promoter was impacted by the introduction of nucleosome positioning elements as well as pre-assembly of nucleosomes using recombinant hypoacetylated wild-type histones or, to promote accessibility, histones containing lysine (K) to glutamine (Q) mutations to mimic acetylated states at histone H3 K9,14 or histone H4 K16 prior to their delivery into cells. By monitoring transient and prolonged expression of the reporter enhanced Green Fluorescent Protein, eGFP, we observed that inclusion of nucleosome positioning-sequences as well as pre-assembly of the reporter plasmid into chromatin prior to delivery into cells impacted the efficiency of transient or prolonged expression of eGFP. Together, our observations indicated that strategies to promote chromatin accessibility by introduction of nucleosome positioning elements adjacent to transgenes in plasmid-based vectors, or pre-assembly of vectors into chromatin containing attributes of active chromatin states, have the potential to enhance transgene expression in human cells.

## 2. Materials and Methods

### 2.1. Cloning of reporter plasmids

The plasmid peGFP, expressing the enhanced Green Fluorescence Protein (eGFP) under the control of the human Elongation Factor 1 Alpha (EF1α) promoter [53,54] (Figure S1A), was used to generate plasmids pW601-eGFP and pW601-100b-eGFP. The E1Fa promoter was chosen as this promoter prolongs transgene expression without affecting vector copy number [55], whereas the commonly used cytomegalovirus promoter (CMVp) is prone to epigenetic silencing [56,57]. To generate pW601-eGFP, an array of four direct repeats of a 177 bp sequence that contained the 147 bp Widom601 sequence[58] was amplified by PCR from plasmid pUC19-Widom601 [59] using primers oALK1723/1724 (5’ GCGAAGCTTGAATTCGAGCTCGGTACCCG 3’ / 5’ GCCAAGCTTGCATGCCTG 3’, Integrated DNA Technologies (IDT)). The amplified Widom601 fragment was then digested with HindIII (New England Biolabs) and gel-isolated using the QIAquick Gel Extraction Kit (50) (Qiagen). Purified products were cloned into the unique HindIII site of peGFP (Figure 1). To generate the pW601-100b-eGFP, a 100 base pair fragment was amplified from pUC19 (#50005, Addgene) (Figure S1B) using primers oALK1765/1766 (5’ GCTTGCATGCCTGCAGGCCTGGGGTGCCTAATGAG 3’/5’ CGTCTTCGATATCCTGCATAATGCAGCTGGCACGAC 3’, IDT), then digested with SbfI (New England Biolabs) and gel-isolated. Isolated products were cloned into the unique SbfI site of pW601-eGFP using NEBuilder® HiFi DNA Assembly Master Mix (#E2621S, New England Biolabs). Plasmids were validated by restriction enzyme digestion using HindIII or SbfI-HF and sequenced using primers oALK1727/oALK1728 (5’ ACCGGTTCAATTGCCGAC 3’/5’ CCACAACTAGAATGCAGTG 3’, IDT). Expression of eGFP was tested by epifluorescence microscopy analyses in transient expression assays. Plasmids were isolated under endotoxin-free conditions using the PureLink HiPure Plasmid DNA Purification Kits (K2100-02, Invitrogen) from DH5α E. coli prior to chromatin assembly.

**Figure 1.**
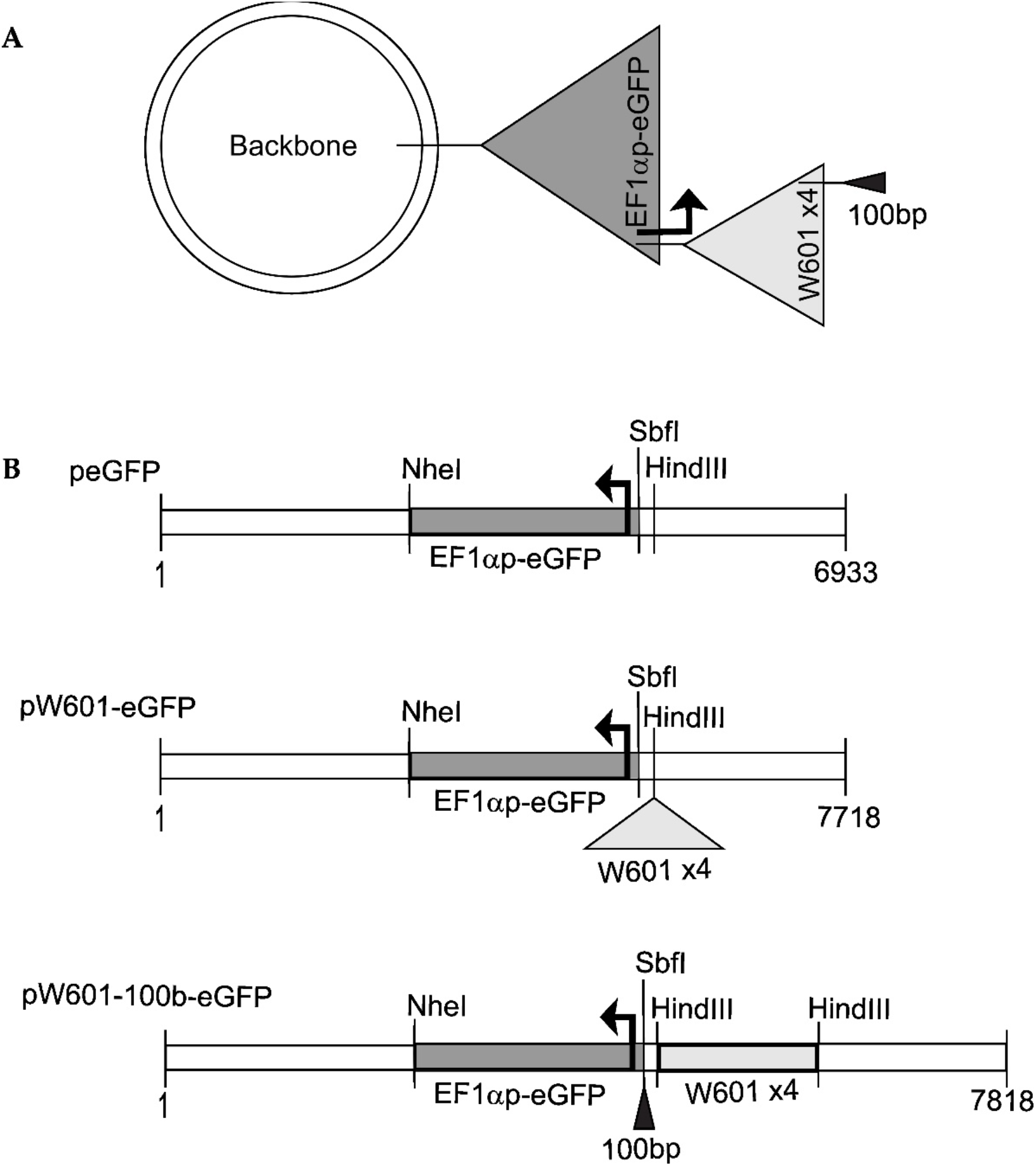
Reporter plasmids used in this study. (**a**) Cloning strategy. A plasmid backbone containing beta-lactamase and aminoglycoside 3’-phosphotransferase (APH (3’)) genes, and bacteria origin of replication (ori) (not shown) was used as the parent plasmid for cloning eGFP reporter plasmids (see Materials and Methods); (**b**) Organization of plasmids peGFP, pW601-eGFP and pW601-100b-eGFP. peGFP contains the eGFP gene expressed from the human EF1α promoter (sequence in Figure S1A). pW601-eGFP contains an array of four 147 bp direct repeats of the Widom601 nucleosome positioning sequence [58] with a 30 bp linker sequence between each repeat, cloned into a HindIII site of peGFP. pW601-100b-eGFP contains a 100 bp fragment cloned into the SbfI site of pW601-eGFP. Genes and other sequences are not drawn to scale.

### 2.2. Site-directed mutagenesis of H3K9,14 and H4K16 to mimic acetylated states

Plasmids pET21b encoding histone H3 and H4 genes [60] from Xenopus laevis were used to generate H3K9,14Q or H4K16Q mutants. The H3K9Q mutant was generated by site-directed mutagenesis using primers 5’ CGCCCGTCAGTCCACCGGAG 3’ / 5’ CCGGTGGACTGACGGGCGG 3’. Then, the H3 K9Q mutant was used to generate the H3K9,14Q mutant similarly using primers 5’ CGTAAATCCACCGGAGGGCAGGCTCCCCGCAAGCAGC 3’ / 5’ GCTGCTTGCGGGGAGCCTGCCCTCCGGTGGATTTACG 3’. The H4K16Q mutant was generated similarly using primers 5’ GGGTAAAGGTGGTGCTCAGCGTCACCGTAAAGTTC 3’ / 5’ GAACTTTACGGTGACGCTGAGCACCACCTTTACCC 3’. Mutated sequences were verified via Sanger sequencing.

### 2.3. Refolding of histone octamers and assembly of chromatin

Recombinant core histones (H2A, H2B, H3, H4), or mutants (H3K9,14Q and H4K16Q) from Xenopus laevis were expressed and purified from E. coli (BL21 (DE3)) as described in our previous studies [59,61]. To assemble plasmids into chromatin while mimicking the length of DNA in a nucleosome [58,62,63], recombinant unmodified histone octamers were mixed with plasmids at a molar ratio of 0.3 moles of histone octamers per every 177 bp DNA length, corresponding to 147 bp nucleosomal DNA plus a 30 bp linker DNA sequence, as cloned in the Widom601 array-containing plasmids. Mixed samples were reconstituted into chromatin under salt gradient dialysis as described in their studies[60,64,65]. Successful assembly was validated by DNA gel electrophoresis, see Figure S2 for representative validation analysis.

### 2.4. Transfection by calcium phosphate

5×10^4^ 143B cells (CRL-8303, ATCC) were seeded per well on a 6-well plate for 24 h prior to transfection. 0.6 picomole of plasmids pre-assembled into chromatin or unassembled plasmid DNA, and a final culture concentration of 11.5 mM CaCl_2_ was used per transfection. Briefly, a 230 µl mixture per transfection was generated by adding 2X HEPES-Buffered Saline (HBS)-pH 7.0 containing 50 mM HEPES, 280 mM NaCl and 1.5 mM Na_2_HPO_4_ to a final concentration of 1X HBS, followed by DNA and lastly, 100 mM CaCl_2_. The mixture was allowed to incubate at room temperature for 10 minutes to allow formation of calcium-phosphate precipitates prior to adding to cell cultures [66]. 24 h post-transfection, cell supernatant was removed, and cells were supplemented with fresh growth media containing DMEM, 10% fetal bovine serum, and 100 units of penicillin-streptomycin. Cultures were maintained in 37 °C incubator containing 5% CO_2_.

### 2.5. Flow cytometry analyses of expression of reporter eGFP

Untransfected 143B cells or 143B cells transfected with plasmids that encoded the reporter eGFP were trypsinized with 1X Trypsin-EDTA (0.5%) and aliquots of cells were stained with 0.4% trypan blue, counted using a hemocytometer, and 1×10^5^ viable cells were isolated by centrifugation at 300xg for 3 min at room temperature. Cells were resuspended in 1X phosphate buffered saline (1X PBS) containing 137 mM NaCl, 2.7 mM KCl, 10 mM Na_2_HPO4, and 1.8 mM KH2PO4, pH 7.4. Cell pellets were resuspended in 200 µl 1X PBS and kept on ice until flow cytometry. Samples were filtered through cell strainer FACS tubes (Stellar Scientific, FSC-9005) to remove clumped cells just prior to flow cytometry analysis. Samples were analyzed using a BD Accuri C6 Plus flow cytometer and BD Accuri C6 Plus software, version 1.0.27.1. The expression of reporter eGFP was analyzed using the FL-1 488 nm channel. Mean fluorescence intensity in the FL-1 488 nm channel below 1×10^4^ was set to background in this study. Untransfected cells served as the negative control. See supplemental Figure S3 for representative epifluorescence microscopy images and example on flow cytometry analysis strategy used herein.

### 2.6. Statistical analysis of expression of reporter eGFP

One-Way Anova [67] was used to compare the individual means of the percent eGFP+ cells from a given plasmid construct to the overall mean of percent eGFP+ cells from all plasmids within an experiment. One-Way Anova was also used to compare mean fluorescence intensity of those cells expressing eGFP from a given plasmid construct to the overall mean of all plasmids tested. To obtain *p* values, Tukey’s Honestly-Significant Difference (TukeyHSD) [68] post-hoc test was used to perform pairwise comparisons of eGFP expression between plasmid constructs. *p* values *< 0.05* were considered significant.

### 2.7. Rate of loss of expression of reporter eGFP

To determine the rate of loss of cells expressing eGFP, or the rate of loss of fluorescence intensity from the eGFP+ subpopulation, the number of cell generations at every time point from Day 3 (D3) to Day 9 (D9) were calculated as the number of hours after transfection divided by the cell doubling time (Equation 1). The doubling time was computed as the number of hours after transfection times log (2) divided by the difference between log of final cell count, at the time of analysis, and log of initial cell count, at the time of seeding (Equation 2). For the rate of loss of percent eGFP+ cells or the fluorescence intensity, the slope (m) of the linear regression line, Y = mX + b, where Y was percent eGFP+ cells or fluorescence intensity, X was the number of cell generations, and b was the Y-intercept (Equation 3) was computed.

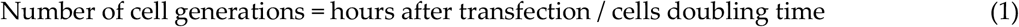

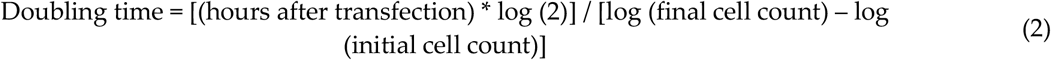

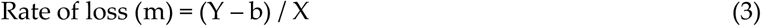

## 3. Results

### 3.1. Cis-sequences impact the efficiency of expressin of eGFP from plasmids transfected as naked DNA

In mammals, ∼147 bp of negatively charged DNA is wrapped around a positively charged histone octamer to form a nucleosome through electrostatic interactions, and adjacent nucleosomes are separated by inter-nucleosomal “linker DNA” within chromatin[69]. Certain DNA sequence motifs such as the artificially derived Widom601 sequence have high affinity for histone octamers and can ‘position’ the octamer at these motifs as well as promote positioning of neighboring nucleosomes into ordered, or phased, arrays upon nucleosome assembly both in vitro and in vivo [58,70-72]. To establish the impact of cis-sequences capable of promoting nucleosome positioning on the efficiency of expression and retention of reporter eGFP, we generated a series of reporter plasmids encoding the eGFP expressed from the human EF1α promoter [53,54,73] (Figure 1, Figure S1). Four 177 bp direct repeats containing the Widom601 sequence [58,70] were cloned upstream the EF1α promoter (Figure 1) to drive assembly of positioned nucleosomes and to promote phasing of nucleosomes across adjacent DNA sequences [74]. The TATA box-containing EF1α promoter has several regulatory regions that facilitate efficient promoter activity [53,54] (Figure S1A). As nucleosome phasing adjacent to nucleosomes positioned by the Widom601 array had the potential to affect accessibility of cis-acting regulatory elements within the EF1α promoter, a third construct containing 100 bp fragment cloned between the Widom601 array and the EF1α promoter was also generated to shift nucleosome phasing across the promoter region.

To assess the impact of nucleosome positioning sequences on the efficiency of expression and retention of reporter eGFP from plasmids transfected as “naked DNA”, the parent plasmid (peGFP), the plasmid containing Widom601 array (pW601-eGFP) or the plasmid containing the Widom601 array plus an 100 bp insert (pW601-100b-eGFP) (Figure 1B) were transfected into human 143B cells and assessed for the efficiency of expression of eGFP at ∼72 h (D3), 144 h (D6) and 216 h (D9) post-transfection by flow cytometry. Expression of eGFP was assessed by two parameters; the percentage of cells in the population that expressed eGFP, as well as the intensity of fluorescence of the eGFP+ subpopulation (Figure 2). eGFP was expressed in 15.8 ± 4.6%, n = 6, of the cells transfected with peGFP at three days post-transfection. This eGFP+ subpopulation had a mean fluorescence intensity of 890,541 ± 68,348, n = 6. Introduction of the Widom601 array upstream of the EF1α promoter (pW601-eGFP) did not significantly impact the percentage of cells transiently expressing eGFP relative to those cells transfected with the parent plasmid peGFP (Figure 2A). In contrast, transfection with pW601-100b-eGFP resulted in a significantly greater percentage of eGFP+ cells relative to that observed for either peGFP or pW601-eGFP at three days post-transfection, *p = 0.0004* and *p = 0.0036*, n = 6, respectively. Similarly, the eGFP+ subpopulation in cells transfected with pW601-100b-eGFP exhibited a higher mean fluorescence intensity relative to that observed in cells transfected with peGFP or pW601-eGFP at three days post-transfection, *p = 0.0245* and *p = 0.0165*, n = 6, respectively (Figure 2B). Together, these observations indicated that sequences capable of influencing nucleosome positioning could alter the percentage of cells initially expressing eGFP as well as the level of eGFP expressed in those cells.

**Figure 2.**
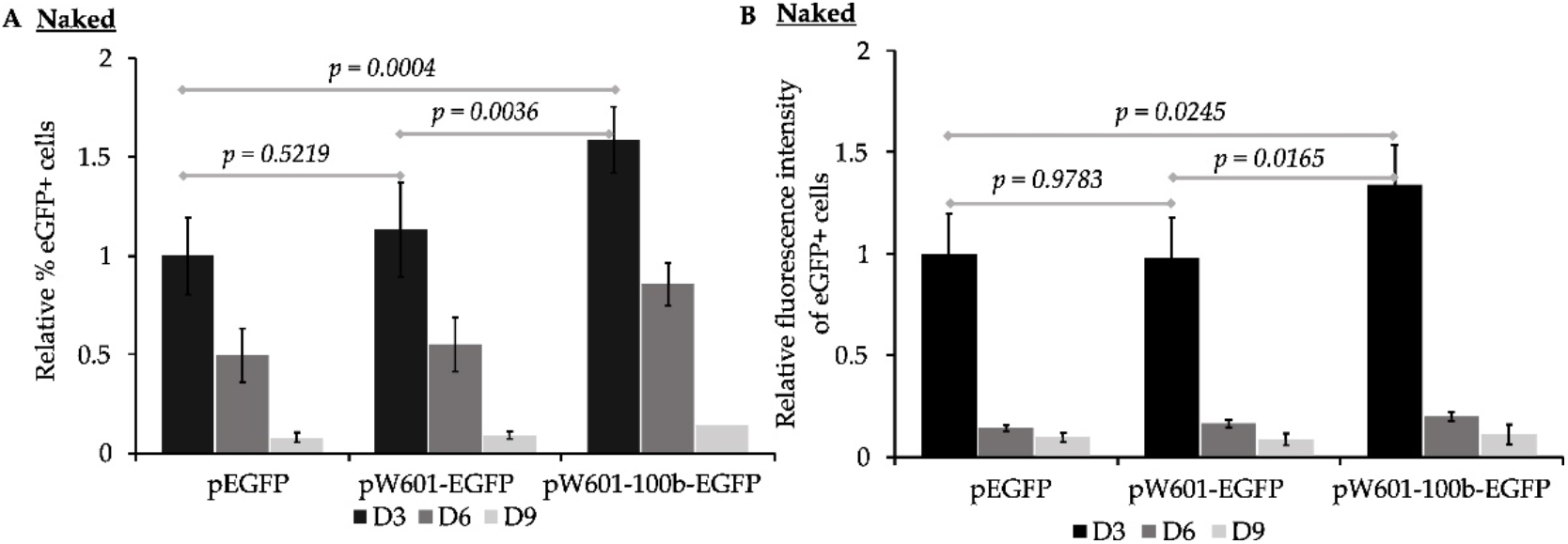
Nucleosome positioning sequences impact the efficiency of expression of eGFP from plasmids lacking pre-assembled nucleosomes. 143B cells transfected with the indicated plasmids (lacking pre-assembled nucleosomes) were analyzed for expression of eGFP by flow cytometry at ∼72 h (D3), ∼144 h (D6), and ∼216 hours (D9) post-transfection. (**a**) Percent eGFP+ cells at D3, D6 and D9. Percent eGFP+ cells were normalized relative to percent eGFP+ cells transfected with peGFP at D3, which was set to 1 (% eGFP+ cells of indicated sample at D3, D6, or D9 /% eGFP+ cells transfected with peGFP at D3); Mean ± STD, n = 6); (**b**) Mean fluorescence intensity of the eGFP+ subpopulations in Figure 2A were determined for each sample and timepoint, and then normalized relative to the mean fluorescence intensity of cells transfected with peGFP at D3, which was set to 1 (Fluorescence intensity of indicated sample at D3, D6, or D9 / Fluorescence intensity of cells transfected with peGFP at D3; Mean ± STD, n = 6). Statistical analyses were conducted using One-way Anova [67], and indicated *p* values were calculated using the TukeyHSD post-hoc test [68].

To assess whether cis-elements capable of promoting nucleosome positioning impacted short-term retention of expression of eGFP, the rate of loss of eGFP+ cells per generation from the transfected populations was determined for cells that had been transfected with plasmids that lacked (peGFP) or contained nucleosome positioning sequences (pW601-eGFP and pW601-100b-eGFP) (Table 1). Over the course of nine days, eGFP+ cells were lost at similar rates from cell populations transfected with either peGFP or pW601-eGFP, whereas cells transfected with pW601-100b-eGFP lost the eGFP+ subpopulation more rapidly relative to those transfected with peGFP, *p = 0.0066*, n = 6 (Table 1). In contrast, no significant difference was observed in the rate of loss of intensity of fluorescence from the eGFP+ subpopulations over time regardless of whether cells had been transfected with plasmids containing or lacking nucleosome positioning sequences (Table 1). Together, these results are consistent with cis-sequences capable of altering nucleosome phasing influencing retention of expression of transgenes over time.

**Table 1.** Loss of expression of eGFP from reporter plasmids lacking pre-assembled nucleosomes per cell generation.

| Unassembled Plasmids | % Loss of eGFP+ Cells/Generation<sup>a</sup> | % Loss of Fluorescence Intensity/ Generation<sup>a</sup> |
| --- | --- | --- |
| peGFP | 18.4 ± 4<sup>b</sup> | 17 ± 4<sup>e</sup> |
| pW601-eGFP | 20.3 ± 5<sup>c</sup> | 15.4 ± 4<sup>f</sup> |
| pW601-100b-eGFP | 25.7 ± 3<sup>d</sup> | 22.2 ± 7.6<sup>g</sup> |

<sup>a</sup>n = 6; <sup>b&c</sup> p = 0.5175; <sup>a&d</sup> p = 0.0066; <sup>c&d</sup> p = 0.0632; <sup>e&f</sup> p = 0.4917; <sup>e&g</sup> p = 0.2039; <sup>f&g</sup> p = 0.1011. Statistical significance was determined using One-way Anova and p values are from TukeyHSD post-hoc test.

### 3.2. Pre-assembled chromatin retain expression of eGFP more efficiently than naked plasmid DNA

Accessibility of regulatory sequences within nucleosomal DNA by transcription machinery can be facilitated via post-translational modification of histones to disrupt charge-charge interactions between the histone tails and DNA. Transcriptionally repressed regions and condensed chromatin are enriched with deacetylated histones [30,31,75,76]. In contrast, acetylation of lysine residues in histone tails, which neutralizes their positive charge, is associated with enhanced accessibility of DNA and gene expression [28,29,77,78], and transcriptionally active regions are enriched with acetylated histones including acetylated histone H3 at K9 [33], and K14 [33,79], and of histone H4 at K16 [29,80]. Consistent with histone acetylation states influencing expression from the EF1α promoter, treatment with either Class I or Class II histone deacetylase inhibitors at time of transgene delivery via plasmid vectors enhances transgene expression from the EF1α promoter [81-83]. To assess the impact of different pre-assembled chromatin states on the efficiency of expression of eGFP, peGFP was first pre-assembled into nucleosomal DNA in vitro using histone octamers that contained either recombinant unmodified histones H2A, H2B, H3 and H4 or histone octamers containing H2A, H2B, H3K9,14Q, and H4 (Figure 3) as the neutral charge of acetylated lysine residues can be mimicked by mutating Lysine (K) to glutamine (Q) [77,78]. “Naked” peGFP or pre-assembled constructs were then transfected into 143B cells, and the percent eGFP+ cells plus the intensity of fluorescence of those eGFP+ cells were evaluated as in Figure 2. eGFP was expressed in 15.7 ± 0.4%, n = 4, of cells transfected with peGFP lacking pre-assembled nucleosomes at three days post transfection. The mean fluorescence intensity of this eGFP+ subpopulation was 430,944 ± 52,314, n = 4. In contrast, cells transfected with peGFP that had been pre-assembled with either H3- or H3K9,14Q-containing histone octamers had fewer eGFP+ cells in the population at three days post-transfection relative to cells transfected with naked peGFP, p < 0.0001 and p < 0.0001, n = 4, respectively (Figure 3A). The percentage of eGFP+ cells in cells transfected with peGFP that had been pre-assembled into chromatin with unmodified H3-containing octamers prior to transfection was greater than those assembled into chromatin with octamers containing H3K9,14Q at three days post-transfection, p = 0.0127, n = 4 (Figure 3C). However, the mean fluorescence intensity of the eGFP+ subpopulations from all conditions tested were similar at three days post-transfection (Figure 3D). Together, these observations indicate that pre-assembled chromatin states could impact the percentage of cells that initially expressed transgenes; pre-assembly of unmodified, “deacetylated” histones reduced initial levels of expression of eGFP, but mimicking the acetylated state on residues 9 and 14 of H3 in this context was insufficient to promote gene expression. However, the assembled chromatin states did not alter the initial level of expression of eGFP within the eGFP+ subpopulations relative to naked peGFP.

**Figure 3.**
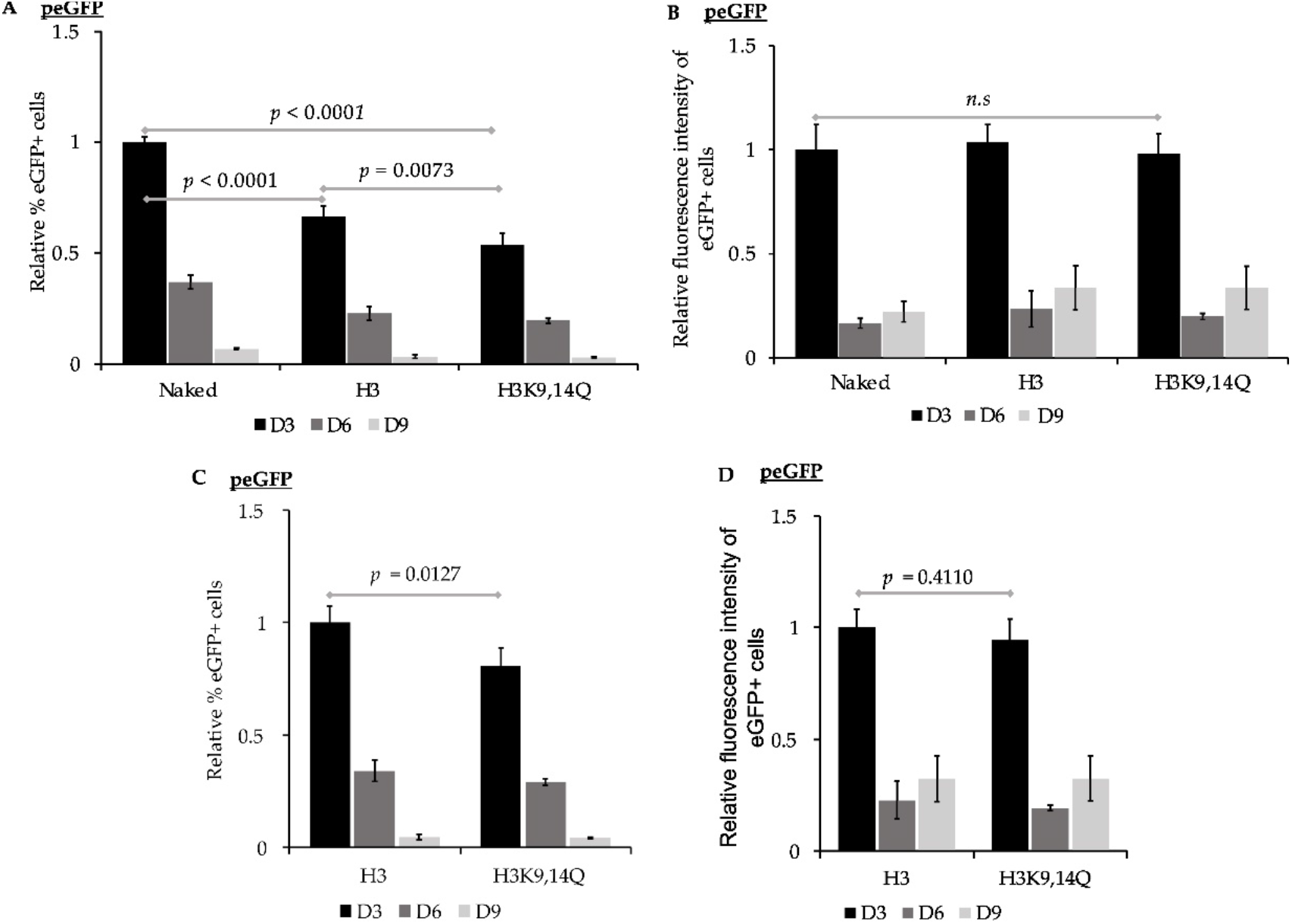
Impacts of pre-assembling peGFP into chromatin on expression of eGFP. 143B cells transfected with peGFP (unassembled), peGFP pre-assembled with unmodified recombinant histone octamers (H3) or with histone octamers containing H3K9,14Q were analyzed for expression of eGFP by flow cytometry at ∼72 h (D3), ∼144 h (D6), and ∼216 hours (D9) post-transfection. (**a**) Percent eGFP+ cells at D3, D6 and D9 were calculated for samples transfected with the indicated constructs, and then normalized relative to percent eGFP+ cells at D3 from cells transfected with (Naked) peGFP, which was set to 1 as in Figure 2 (Mean ± STD, n = 4); (**b**) Mean fluorescence intensity of eGFP+ subpopulations in Figure 3A were calculated and then normalized relative to the mean fluorescence intensity at D3 of cells transfected with (Naked) peGFP, which was set to 1 (see Figure 2, Mean ± STD, n = 4); (**c&d**) Impact of H3 vs. H3K9,14Q-containing chromatin on expression of eGFP. (c) Percent eGFP+ cells at D3, D6 and D9 from cells transfected with the peGFP pre-assembled with histone octamers containing H3 (H3) or H3K9,14Q (H3K9,14Q) were calculated and then normalized relative to percent eGFP+ cells at D3 from cells transfected with peGFP pre-assembled with unmodified histones (H3), which was set to 1 (Mean ± STD, n = 4). (d) Mean fluorescence intensity of eGFP+ subpopulation in Figure 3C was normalized relative to mean fluorescence intensity of eGFP+ subpopulation at D3 from cells transfected with peGFP pre-assembled with unmodified histone octamers (H3), which was set to 1 (Mean ± STD, n = 4). Statistical significance and *p* values were determined as in Figure 2.

Next, we determined the rate of loss of expression of eGFP in cells transfected with naked peGFP or with peGFP pre-assembled into chromatin containing either unmodified H3 or H3K9,14Q (Figure 3 and Table 2). Cells transfected with peGFP pre-assembled into unmodified nucleosomes retained expression of eGFP more efficiently than those transfected with naked peGFP, *p = 0.0012*, n = 4 (Table 2). Similarly, cells transfected with peGFP pre-assembled into nucleosomes containing H3K9,14Q retained expression of eGFP more efficiently than naked peGFP, *p = 0.0008*, n = 4, or peGFP assembled into unmodified nucleosomes, *p = 0.0141*, n = 4. In contrast, no differences in retention of the intensity of eGFP fluorescence signal over time were observed in those eGFP+ subpopulations across all constructs (Table 2). Together, these observations imply that delivery of reporter plasmids pre-assembled into different forms of chromatin can influence both the initial and continued expression of transgenes relative to naked gene delivery.

**Table 2.** Loss of expression of eGFP from peGFP pre-assembled into chromatin containing unmodified histones, or H3K9,14Q acetyl-mimics.

| Chromatin State of peGFP | % Loss of eGFP+ Cells/ Generation<sup>a</sup> | % Loss of Fluorescence Intensity/ Generation<sup>a</sup> |
| --- | --- | --- |
| Unassembled | 10.6 ± 0<sup>b</sup> | 9.4 ± 1.3<sup>e</sup> |
| H3 | 7.7 ± 1<sup>c</sup> | 9.1 ± 1.8<sup>f</sup> |
| H3K9,14Q | 6.0 ± 0<sup>d</sup> | 8.3 ± 1.8<sup>g</sup> |

<sup>a</sup> $n = 4$ ; <sup>b&c</sup> $p = 0.0012$ ; <sup>b&d</sup> $p = 0.0008$ ; <sup>c&d</sup> $p = 0.0141$ ; <sup>e&f</sup> $p = 0.7808$ ; <sup>e&g</sup> $p = 0.3697$ ; <sup>f&g</sup> $p = 0.5706$ . Statistical significance and $p$ values were determined as in Table 1.

### 3.3. Nucleosome positioning elements affect the efficiency of expression of reporter eGFP in pre-assembled chromatin states

Nucleosome positioning can influence the expression state of endogenous genes [69,84-86]. To assess the impact of nucleosome positioning on expression of transgenes from plasmids that had been pre-assembled into chromatin prior to delivery into cells, peGFP, pW601-eGFP, or pW601-100b-eGFP were initially pre-assembled into nucleosomes containing recombinant unmodified histones, then transiently transfected into 143B cells. The efficiency of expression of eGFP was evaluated by determining both the percent eGFP+ cells as well as the intensity of eGFP fluorescence in the eGFP+ subpopulation (Figure 4A&B) as above. When cells were transfected with unmodified nucleosomal peGFP, eGFP was expressed in 23.8 ± 3% of the cells at three days post-transfection, and this eGFP+ subpopulation had a mean fluorescence intensity of 402,776 ± 44,433, n = 4. Introduction of the Widom601 array upstream of the transcriptional start site at the EF1α promoter (pW601-eGFP) reduced the percentage of eGFP+ cells at three days post-transfection relative to cells transfected with unmodified nucleosomal peGFP, *p < 0.0001*, n = 4 (Figure 4A). In contrast, insertion of the 100 bp fragment between the Widom601 array and the EF1α promoter to shift nucleosome phasing (pW601-100b-eGFP) partially suppressed the Widom601-dependent defects (pW601-eGFP) in percent eGFP+ cells at three days post-transfection, *p = 0.0003*, n = 4 (Figure 4A). However, the percentage of eGFP+ cells was not fully restored to levels observed for cells transfected with unmodified nucleosomal peGFP at three days post-transfection, *p = 0.0254*, n = 4 (Figure 4A). Inclusion of the Widom array (pW601-eGFP) similarly reduced the mean fluorescence intensity of the eGFP+ subpopulation at three days post-transfection relative to either peGFP or pW601-100b-eGFP, *p < 0.0001* and *p < 0.0001*, n = 4, respectively (Figure 4B). In contrast, the mean fluorescence intensity of the eGFP+ subpopulations from cells transfected with peGFP or pW601-100b-eGFP were similar (Figure 4B). When comparing retention of eGFP expression as a function of time from plasmids that had been pre-assembled into nucleosomes with unmodified histones, cells that expressed eGFP from pW601-eGFP retained expression more efficiently than did cells transfected with peGFP or pW601-100b-eGFP, *p = 0.0004* and p = *0.0002*, n = 4, respectively (Table 3). Similarly, the eGFP+ subpopulation from pW601-eGFP retained the intensity of eGFP fluorescence more efficiently over time compared to those eGFP+ subpopulations from peGFP- and pW601-100b-eGFP-transfected cells, *p = 0.0015* and *p = 0.0005*, respectively (Table 3). These observations highlight impacts of nucleosome positioning in influencing expression of transgenes from the E1Fα promoter.

**Table 3.** Loss of expression of eGFP from reporter plasmids pre-assembled into chromatin containing unmodified histones.

| Pre-assembled with Unmodified Histones | % Loss of eGFP+ Cells/Generation<sup>a</sup> | % Loss of eGFP Fluorescence/Generation<sup>a</sup> |
| --- | --- | --- |
| peGFP | 11.3 ± 1<sup>b</sup> | 9.1 ± 2<sup>e</sup> |
| pW601-eGFP | 5.3 ± 0<sup>c</sup> | 2.0 ± 1<sup>f</sup> |
| pW601-100b-eGFP | 9.8 ± 0<sup>d</sup> | 9.5 ± 0<sup>g</sup> |

<sup>a</sup> $n = 4$ ; <sup>b&c</sup> $p = 0.0004$ ; <sup>b&d</sup> $p = 0.0928$ ; <sup>c&d</sup> $p = 0.0002$ ; <sup>e&f</sup> $p = 0.0015$ ; <sup>e&g</sup> $p = 0.6850$ ; <sup>f&g</sup> $p = 0.0005$ . Statistical significance and $p$ values were determined in Table 1.

**Figure 4.**
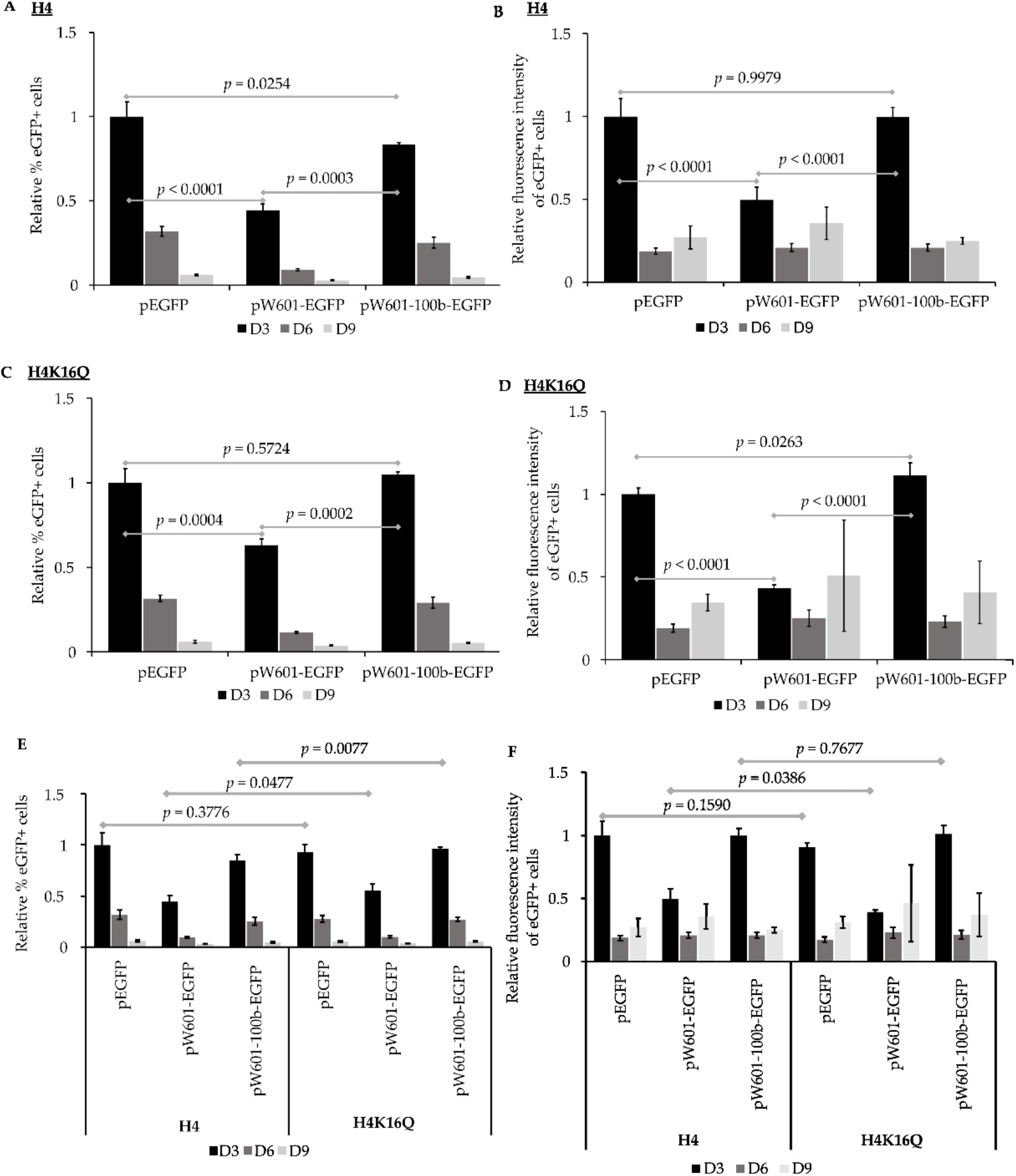
The impact of nucleosome positioning and pre-assembled chromatin states on expression of eGFP. 143B cells transfected with indicated plasmids pre-assembled with unmodified recombinant histone octamers (H4) or pre-assembled with histones octamers containing H4K16Q were analyzed for expression of eGFP by flow cytometry at ∼72 h (D3), ∼144 h (D6), and ∼216 hours (D9) post-transfection. (**a**) Percent eGFP+ cells at D3, D6 and D9 were calculated for samples transfected with the indicated plasmids pre-assembled with unmodified histone octamers (H4). Percent eGFP+ cells was then normalized relative to percent eGFP+ cells at D3 from cells transfected with peGFP pre-assembled with unmodified histone octamers, which was set to 1 (Mean ± STD, n = 4) as in Figure 2; (**b**) Mean fluorescence intensity of eGFP+ subpopulation in Figure 4A was calculated and then normalized relative to mean fluorescence intensity at D3 from cells transfected with peGFP pre-assembled with unmodified histone octamers, which was set to 1 (Mean ± STD, n = 4) as in Figure 2; (**c**) Percent eGFP+ cells at D3, D6 and D9 was calculated for samples transfected with the indicated plasmids pre-assembled with histone octamers containing H4K16Q. Percent eGFP+ cells was then normalized relative to percent eGFP+ cells at D3 from cells transfected with peGFP pre-assembled with H4K16Q histone octamers, which was set to 1 (Mean ± STD, n = 4); (**d**) Mean fluorescence intensity of eGFP+ subpopulation in Figure 4C was calculated and then normalized relative to mean fluorescence intensity at D3 from cells transfected with peGFP pre-assembled with H4K16Q, which was set to 1 (Mean ± STD, n = 4); (**e&f**) Impacts of *cis-*elements and pre-assembled chromatin states on expression of eGFP. (e) Percent eGFP+ cells at D3, D6 and D9 from cells transfected with the indicated plasmids pre-assembled with unmodified histone octamers (H4) or those containing H4K16Q (H4K16Q) (from Figure 4A&C) were re-analyzed by normalizing relative to percent eGFP+ cells at D3 from cells transfected with peGFP pre-assembled with unmodified histone octamers (H4), which was set to 1 (Mean ± STD, n = 4). (f) Mean fluorescence intensities of each eGFP+ subpopulation (from Figure 4B&D) were re-analyzed by normalizing relative to mean fluorescence intensity of the eGFP+ subpopulation at D3 from cells transfected with peGFP pre-assembled with unmodified histone octamers (H4), which was set to 1 (Mean ± STD, n = 4). Statistical significance and *p* values were determined as described in Figure 2.

The acetylation state of histone H4 K16 influences chromatin structure and the acetylated form of this residue promotes the formation of ‘relaxed’ chromatin [29]. Consistent with facilitating access to *cis*-sequence in DNA, H4 K16 acetylation is enriched at enhancers and transcriptional start sites of active genes [87]. To assess the impact of H4 K16 acetylation, or H4 K16 acetylation plus nucleosome positioning on the efficiency of expression of eGFP, histone octamers containing the acetyl mimic H4K16Q were also used to pre-assemble nucleosomes onto peGFP, pW601-eGFP and pW601-100b-eGFP (Figure 4C&D) as part of the experiment shown in Figure 4A and 4B). For 143B cells transfected with peGFP pre-assembled into H4K16Q-containing chromatin, GFP was expressed in 22.2 ± 1.6% of the cells at three days post-transfection. This eGFP+ subpopulation had a mean fluorescence intensity of 365,296 ± 14,127, n = 4. For plasmids pre-assembled into H4K16Q-containing chromatin, fewer eGFP+ cells were observed in 143B cells transfected with pW601-eGFP relative to either peGFP or pW601-100b-eGFP at three days post-transfection, *p = 0.0004* or *p = 0.0002*, n = 4, respectively, whereas percentage of eGFP+ cells was similar between peGFP and pW601-100b-eGFP (Figure 4C). Similarly, the presence of the Widom601 array in pW601-eGFP reduced the MFI of the eGFP+ subpopulation relative to that observed in cells transfected with peGFP or pW601-100b-eGFP at three days post-transfection, *p < 0.0001* and *p < 0.0001*, n = 4, respectively (Figure 4D). Further, the mean fluorescence intensity of this eGFP+ subpopulation was higher in cells transfected with pW601-100b-eGFP compared to peGFP at three days post-transfection, *p = 0.0263* (Figure 4D). Together, these observations are consistent with nucleosome positioning also affecting expression from the E1Fa promoter when H4 K1Q nucleosomes are pre-assembled onto the reporter plasmids.

The data from this experiment (Figure 4A-D) was then reanalyzed to assess whether pre-packaging into nucleosomes containing H4K16Q would facilitate expression of transgenes more efficiently than unmodified histones (Figure 4E&F and Figure S4). In this case, the samples were normalized relative to percent eGFP+ cells from cells transfected with peGFP pre-assembled with H4 (Figure 4E). Similarly, the mean fluorescence intensity of the eGFP+ subpopulations were normalized relative to the mean fluorescence intensity of the eGFP+ subpopulation from cells transfected with peGFP pre-assembled with H4 (Figure 4F). No difference in the percent eGFP+ cells was observed for peGFP pre-assembled with histone octamers containing H4 relative to H4K16Q at three days post-transfection (Figure 4E and Figure S4A&D). In contrast, cells transfected with pW601-eGFP or pW601-100b-eGFP pre-assembled into H4K16Q-containing nucleosomes had a higher percentage of eGFP+ cells relative to those assembled into H4-containing nucleosomes, *p = 0.0477* and *p = 0.0077*, respectively, n = 4, at three days post-transfection (Figure 4E). Despite increasing the percentage of cells that initially expressed eGFP under certain conditions, pre-assembly into H4 K16Q relative to H4-containing nucleosomes did not enhance the intensity of eGFP fluorescence from any of the plasmids at three days post-transfection (Figure 4F and Figure S4B,D&F). Similar to pre-assembly with H4-containing nucleosomes, cells transfected with pW601-eGFP pre-assembled into H4K16Q-containing nucleosomes retained eGFP+ cells in the population over time more efficiently than did cells transfected with similarly pre-assembled peGFP or pW601-100b-eGFP plasmids, *p = 0.0005* and *p = 0.0001*, respectively (Table 4). Likewise, the eGFP+ subpopulation from pW601-eGFP pre-assembled into H4K16Q-containing nucleosomes retained the intensity of eGFP fluorescence over time more efficiently than similarly pre-assembled peGFP or pW601-100b-eGFP plasmids, *p = 0.0155* or *p = 0.0082*, n = 4, respectively (Table 4). However, for all conditions, the fluorescence intensity of the eGFP+ cells initially decreased rapidly between three and six days post-transfection, then stabilized in the eGFP+ cells during later timepoints (Tables 1-4). Thus, these data are also consistent with the Widom601 array influencing nucleosome assembly or positioning during pre-assembly in a manner that limited the maximal fluorescence intensity achieved in the eGFP+ cells at three days post-transfection.

**Table 4.**
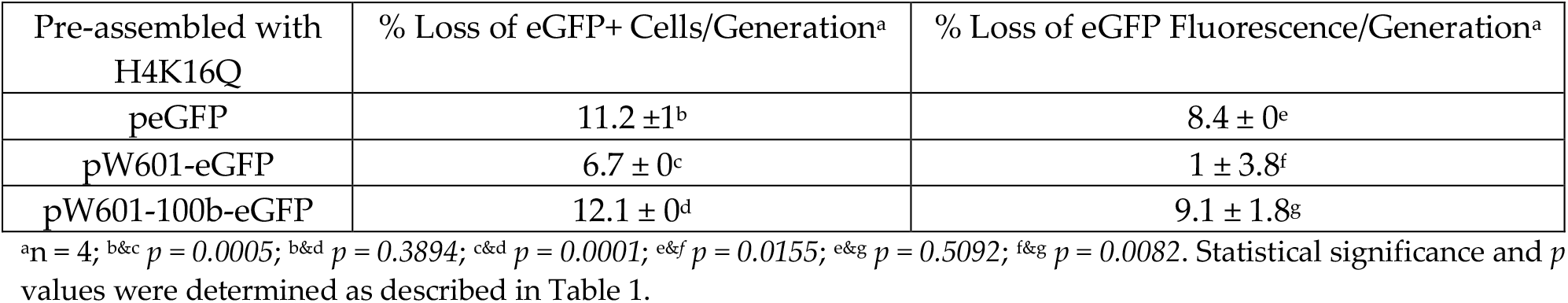
Loss of expression of eGFP from reporter plasmids pre-assembled into chromatin containing H4K16Q.

## 4. Discussion

When naked plasmid DNA is delivered into cells, that DNA becomes assembled into chromatin by the host cell, and this can exert repressive or active effects on gene expression [8,20,21,88]. This study evaluated chromatin accessibility-based strategies, including nucleosome positioning and histone acetylation, for their application in establishing epigenetically transcriptionally active states expression via “priming” the initial characteristics of chromatin at a transgene upon or prior to introduction into cells. Our observations indicated that an array of four Widom 601 nucleosome positioning elements [58,71] could negatively or positively influence short-term expression of transgenes from plasmids in a manner dependent on their distance from EF1α promoter when those plasmids were introduced into cells as naked DNA (Figure 2) or as pre-assembled chromatin (Figure 4). These observations were consistent with host cell-mediated packaging and phasing of nucleosomes on the reporter plasmids upon nuclear entry, as well as pre-assembly of chromatin states, having impacted accessibility of transcription machinery to the EFFa promoter to promote gene expression.

The EF1α promoter contains several regulatory sequences, including binding sites for transcription factors Sp1 [89], Ap1 [90], EFP1 and EFP2 [54] (Figure S1), whose loss can negatively affect gene expression [53,54]. As shifting the distance between the EF1α promoter and the Widom 601 elements by 100 bp promoted initial expression of eGFP, the initial positioning of nucleosomes across the EF1α promoter by the host cell may have influenced transcription (Figure 3 and 4) in a manner affecting *cis*-element accessibility or akin to position-effect variegation [91]. The observation that pW601-100b-eGFP exhibited a more rapid loss of eGFP+ cells over time relative to peGFP (Tables 3 and 4), implied that, while nucleosome positioning could enhance initial transgene expression, nucleosome positioning could also impact the stability of expression over time. Consistent with this, initial high expression of transgenes is often associated with subsequent silencing, perhaps as a result of elevated cellular responses against expression of foreign DNA [12,15].

Efficient gene expression is also facilitated by chromatin post-translational modifications, such as histone acetylation, that influence access to cis-acting regulatory elements by the transcription machinery at promoters [24,25,31]. In this study, pre-assembly of peGFP with histone octamers containing either unmodified recombinant histones or H3K9,14Q mutants reduced the percentage of cells expressing eGFP at three days post-transfection relative to cells transfected with naked peGFP plasmid (Figure 3A). The effects of unmodified histones were consistent with “priming” of the reporter via pre-assembly with hypoacetylated histone octamers having promoted a chromatin state incompatible with efficient gene expression. However, as acetylated H3K9 and H3K14 are enriched at promoters, and correlate with activation of gene expression by RNA Polymerase II [32,33,92], it was somewhat surprising that “priming” with a hyperacetylated state at these residues on histone H3 was insufficient to promote efficient production of eGFP+ cells. These differences in percent eGFP+ cells were not a result of cytotoxic effects of pre-assembled chromatin as we observed no significant differences in cell growth or in the number of trypan blue-positive apoptotic cells [93] (data not shown). Whether these differences also reflect difficulty in cellular uptake of reporter plasmids that are pre-assembled into chromatin versus naked DNA by calcium phosphate-based transfection [66] and/or defects in delivery of the transgene vectors into the nucleus is currently unclear. Future optimization of delivery strategies for chromatinized vectors in the future is warranted to clarify this issue.

While pre-assembled chromatin containing unmodified H3 or H3K9,14ac reduced the percentage of cells initially expressing eGFP, this effect was independent of the mean fluorescence intensity of eGFP observed within those eGFP+ cells (Figure 3). These observations were consistent with, on a cell-to-cell basis, pre-assembled H3 or H3 K9,14ac chromatin having adversely affected an early event such as nuclear entry or initial recruitment of transcriptional machinery to the promoter rather than the expression level that could be achieved. As nuclear delivery of exogenous macromolecules via the nuclear pore is size-selective [94] and large molecules and proteins can be restricted to the cytosol until the nuclear envelope disassembles during mitosis [95,96], such factors may have influenced the efficiency of nuclear delivery of chromatinized plasmids in this study. Also consistent with an nuclear import defect, H3K9,14Q lies in a region used by the nuclear import receptor importin-4 to recognize unmodified H3 [97]. In the absence of nucleosome positioning (Widom601) sequences, in vitro nucleosome assembly likely occurred in a random manner that may have hindered the accessibility of cis elements by transcription factors at the EF1α promoter (Figure S1A), thereby reducing the percentage of cells expressing eGFP (Figure 3). However, cells transfected with plasmids pre-assembled into H3K9,14Q-containing chromatin retained expression of eGFP over time more efficiently than those transfected with naked plasmid DNA or plasmids pre-assembled with hypoacetylated histones (Table 2). Thus, pre-assembled acetylated states on H3 K9 and 14 promoted prolonged expression of eGFP.

When evaluating combined impacts of Widom601 nucleosome positioning elements and pre-assembly of chromatin on plasmids prior to their introduction into cells, distance-specific effects of the Widom601 sequences on the expression of eGFP were observed. The presence of the Widom601 repeats led to reduced expression of eGFP as measured by either the percent eGFP+ cells, or the fluorescence intensity of those eGFP+ cells when the reporter plasmid was pre-assembled into nucleosomal DNA containing unmodified histone octamers, relative to the absence of pre-assembly (Figure 4, Figure 2). This may have reflected nucleosome being positioned over regulatory elements at the promoter during assembly as these effects could be suppressed by altering the distance between the Widom601 sequences and the promoter. Like unmodified H4, pre-assembly with H4K16Q led to a Widom601-dependent and distance-specific reduction in the percent eGFP+ cells and the efficiency of expression of eGFP in the eGFP+ cells (Figure 4). Such Widom601-derived effects on expression of eGFP persisted even when the reporter was pre-assembled into a relaxed chromatin state (H4K16Q), despite H4K16ac being known to promote the formation of relaxed open or accessible DNA [28,29]. However, compared to chromatin containing unmodified H4, “priming” with H4K16Q-containing chromatin could mildly enhance eGFP expression (Figure 4E&F). Together, nucleosome positioning played the predominant role in influencing expression from the EF1α promoter under the conditions tested.

In summary, our findings demonstrated that cis-sequences and pre-assembled chromatin states influenced initial expression and retention of transgenes, and provide a foundation for further development of chromatin-based strategies to increase the probability of forming epigenetically active states of expression upon delivery of transgenes into target cells. The ability to modulate nucleosome positioning and chromatin post-translational modifications have the potential to provide a powerful toolkit for multiple potential applications, including gene therapy, where precise control over gene expression is critical. Future studies assessing impacts of additional modifications, either singly or in combinations, together with cis-acting elements should further refine the control of expression of transgenes. Circumventing potential difficulties in nuclear delivery of pre-assembled chromatin relative to naked DNA may be possible via introducing nuclear targeting sequences to facilitate their nuclear entry [48-50]. Further, combining the chromatin-based strategy described herein with other approaches, including addition of non-B DNA elements [98-100] or Matrix Attachment Regions [101] may also facilitate optimizing expression of nonviral plasmid-based transgenes in the future.

## Supplementary Materials

Figure S1: The human EF1α promoter driving expression of the eGFP reporter gene; Figure S2: Representative analysis of unassembled plasmid DNA or plasmids pre-assembled with unmodified histone octamers; Figure S3: Representative fluorescence microscopy images and representative flow cytometry plots; Figure S4: The impact of pre-assembled nucleosomes containing H4K16Q relative to unmodified histones (H4) within plasmid constructs.

## Author Contributions

Conceptualization, A.L.K. and C.Y.; Methodology, A.L.K. and C.Y.; Investigation, Formal Analysis, and Validation, R.K., J.X. and N.N.; Resources, A.L.K and C.Y.; Writing – Original Draft Preparation, Review & Editing, R.K., J.X., A.L.K. and C.Y.; Visualization, R.K. and J.X.; Supervision and Project Administration, A.L.K. and C.Y.; Funding Acquisition, A.L.K. and C.Y.

## Institutional Review Board Statement

Not applicable.

## Informed Consent Statement

Not applicable.

## Data Availability Statement

The original contributions presented in this study are included in the article. Materials are available upon request.

## Acknowledgments

We thank Bill Sugden for 143B cells. We thank Sihui Wang, Zeyu Zhang, and Arman Sabbaghi at Purdue Statistical Consulting Service for statistical assistance as well as Kyle Wettshurack for technical assistance.

---

**Figure S1.**
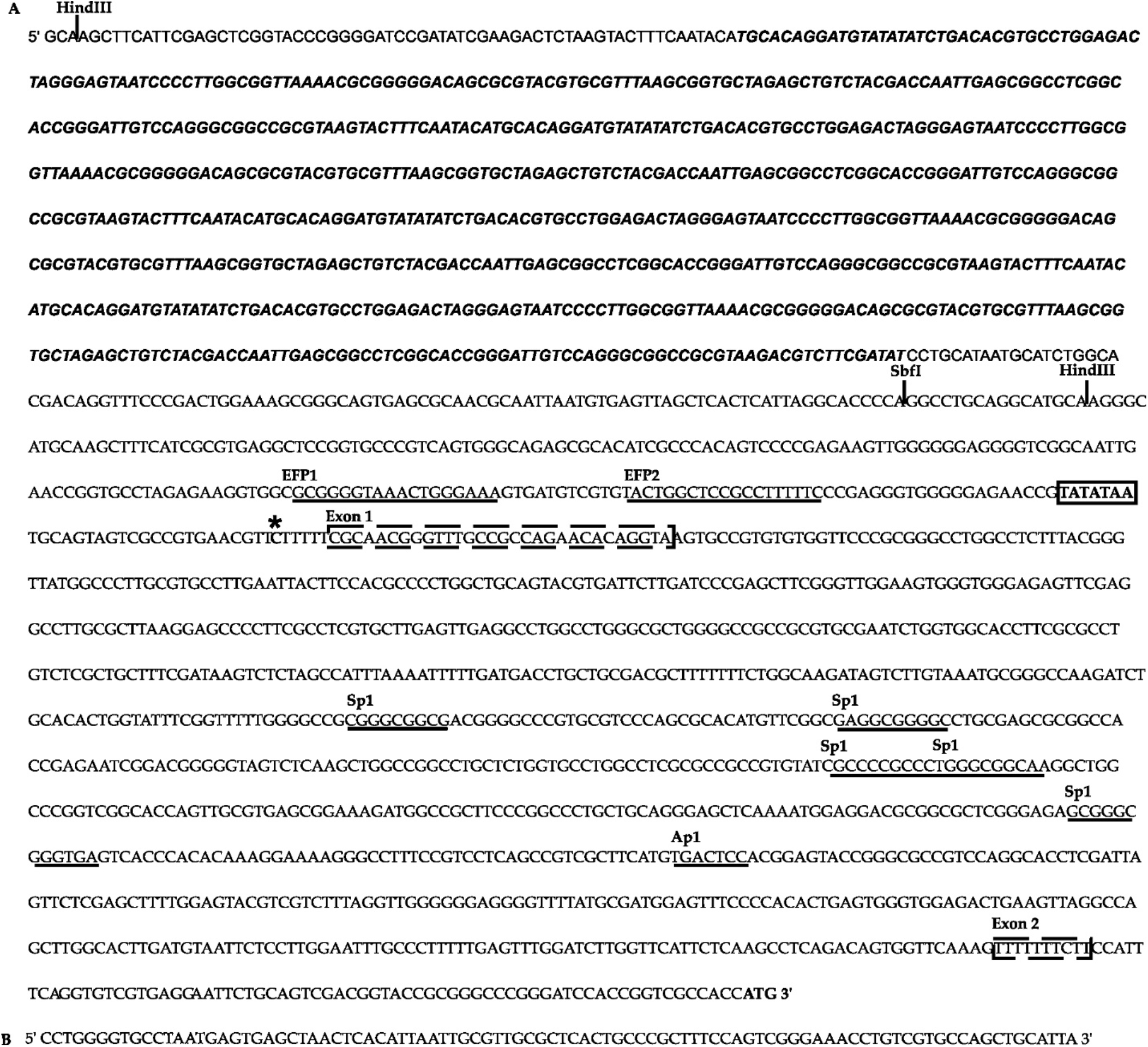
The human EF1α promoter driving expression of the eGFP reporter gene. (**a**) The array of four direct repeats of the 177 Widom601 nucleosome positioning sequence [1,2] in pW601-eGFP plasmid is shown in bold italics. The TATA box is in a solid box. The two untranslated exons (Exon 1 and Exon 2) separated by Intron-A of the EF1α promoter are shown in doKed boxes. Binding sites for transcription factors (EFP1, EFP2, Sp1 and Ap1) are noted and underlined [3,4]. The transcription start site is shown by asterisk. The translation start site of eGFP is noted at the 3’ end of the sequence (ATG in bold); (**b**) Sequence of the 100 bp fragment amplified from pUC19 DNA and cloned into the SbfI site between the Widom601 sequence and the EF1α promoter in plasmid pW601-100b-eGFP.

**Figure S2.**
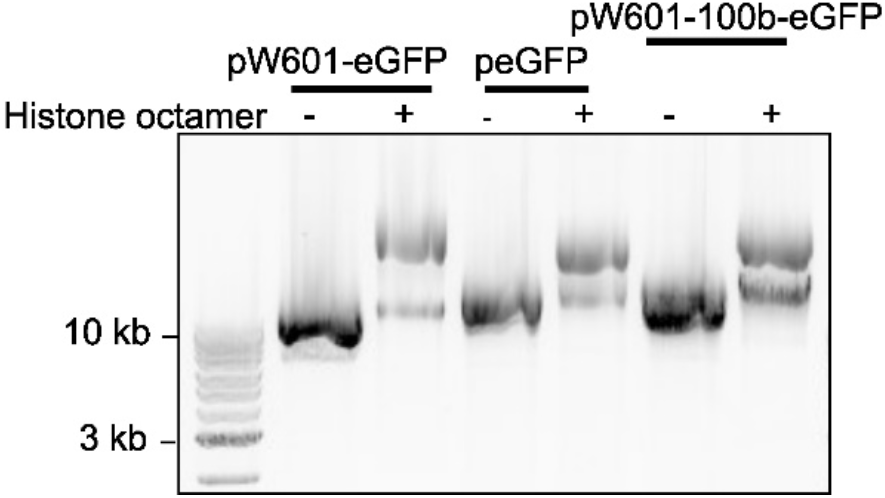
Representative analysis of unassembled plasmid DNA or plasmids pre-assembled with unmodified histone octamers. Samples were evaluated on a 0.8% agarose gel in TBE buffer containing 89 mM Tris base, 89 mM Borate and 2 mM Na_2_EDTA. Chromatin assembly was performed as described in our previous studies [5,6].

**Figure S3.**
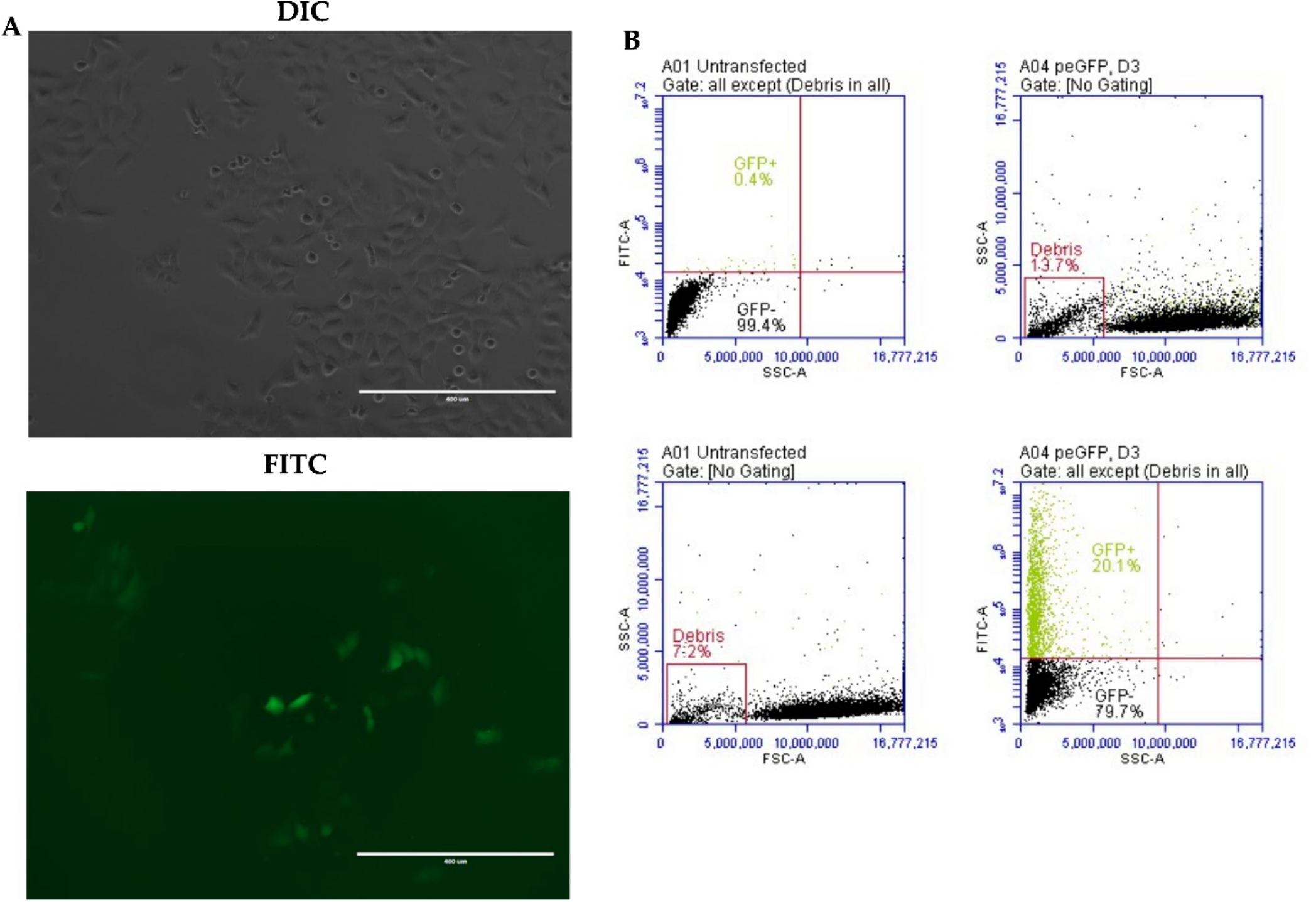
Representative fluorescence microscopy images and representative flow cytometry plots. (**a**) 143B cells transfected with unassembled peGFP were imaged at 72 h post-transfection under sterile live cell-imaging conditions using the EVOS FL inverted fluorescence microscope (Life Technologies). Images were taken at 10X magnification. Scale bar equals 400 µm; (**b**) Untransfected 143B cells (left panels) or 143B cells transfected with unassembled peGFP (right panels) were analyzed by flow cytometry at 72 h post-transfection. Cells were trypsinized and 1×10^5^ viable cells were isolated, washed in 1X PBS, and resuspended in 200 µl 1X PBS as described previously. Side scaKer (SCC-A) and forward scaKer (FSC-A) data were used to assess cell size and complexity. After thresholding to remove “Debris” (top panels) ten thousand events were analyzed, and eGFP+ cells were detected through the FL-1 488 nm channel (FITC-A) against the side scaKer (SCC-A). Mean fluorescence intensity of 1×10^4^ was set as the baseline for the FITC-A channel. Untransfected control cells had 0.4% eGFP+ cells above baseline, while those cells transfected with peGFP had a 20.1% eGFP+ cells at three days post-transfection. The fluorescence intensity of eGFP above baseline ranged from 1×10^4^ to 1×10^7^ (arbitrary units)_(Figure S3B, boKom panels), and was used to calculate the mean intensity of fluorescence signal from the eGFP+ cells (see Materials and Methods).

**Figure S4:**
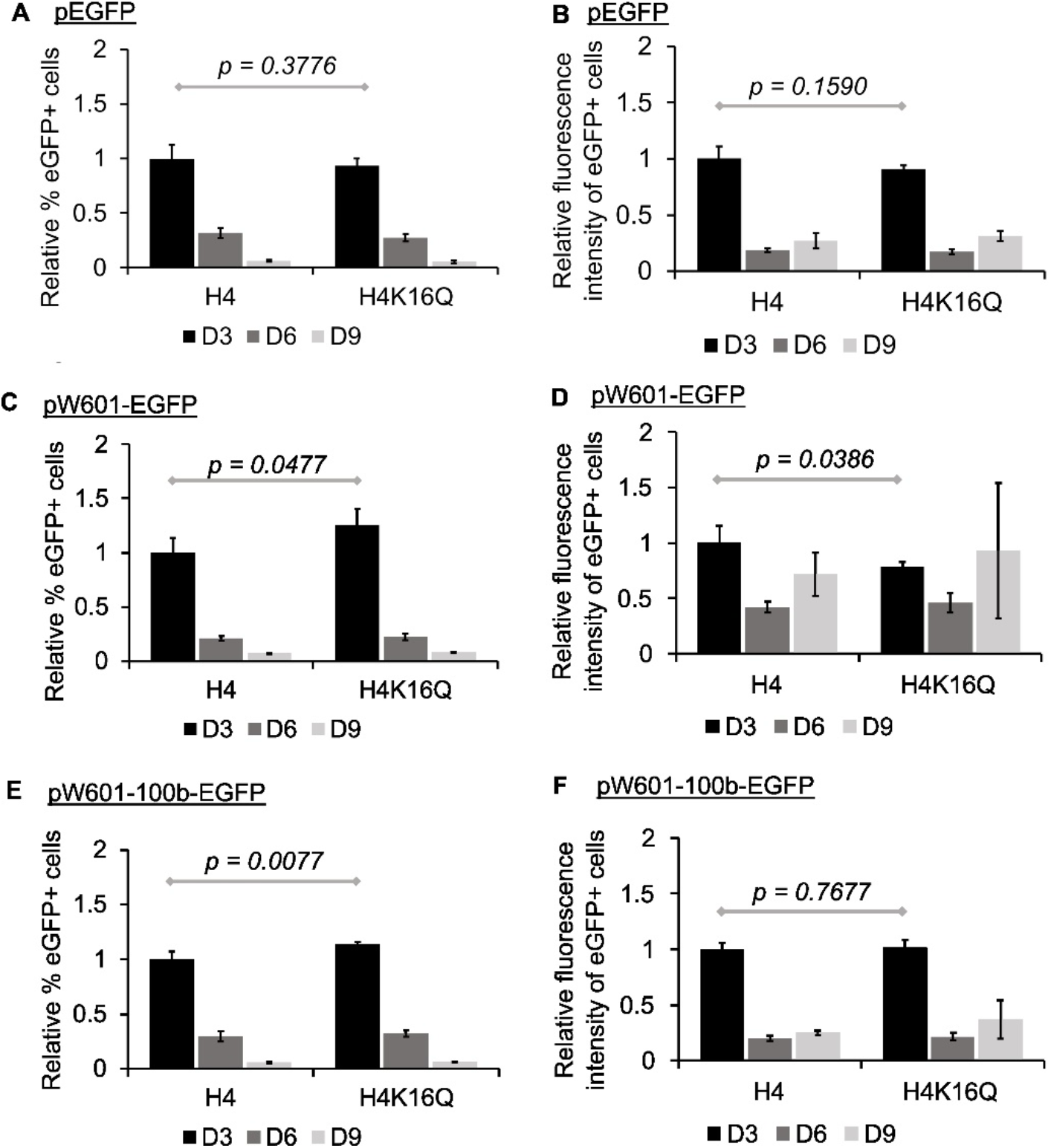
The impact of pre-assembled nucleosomes containing H4K16Q relative to unmodified histones (H4) within plasmid constructs. Re-analysis of data from experiment shown in Figure 4. (**a**,**c&e**) Percent eGFP+ cells from H4K16Q-containing chromatin normalized relative to percent eGFP+ cells from H4-containing chromatin on the indicated plasmids; (**b**,**d&f**) MFI of the eGFP+ subpopulation from H4K16Q-containing chromatin normalized relative to MFI of the eGFP+ subpopulation from H4-containing chromatin on the indicated plasmids. Statistical significance was determined using One-way Anova [7] and *p* values are from TukeyHSD post-hoc test [8].

## References

1. Glover, D.J.; Lipps, H.J.; Jans, D.A. Towards safe, non-viral therapeutic gene expression in humans. Nat. Rev. Genet. 2005, 6, 299–310, doi:10.1038/nrg1577.

2. Naso, M.F.; Tomkowicz, B.; Perry, W.L., 3rd; Strohl, W.R. Adeno-Associated Virus (AAV) as a Vector for Gene Therapy. BioDrugs 2017, 31, 317–334, doi:10.1007/s40259-017-0234-5.

3. Berns, K.I.; Muzyczka, N. AAV: An Overview of Unanswered Questions. Hum. Gene. Ther. 2017, 28, 308–313, doi:10.1089/hum.2017.048.

4. Gonçalves, G.A.R.; Paiva, R.M.A. Gene therapy: advances, challenges and perspectives. Einstein 2017, 15, 369–375, doi:10.1590/s1679-45082017rb4024.

5. Shchaslyvyi, A.Y.; Antonenko, S.V.; Tesliuk, M.G.; Telegeev, G.D. Current State of Human Gene Therapy: Approved Products and Vectors. Pharmaceuticals 2023, 16, doi:10.3390/ph16101416.

6. Bulcha, J.T.; Wang, Y.; Ma, H.; Tai, P.W.L.; Gao, G. Viral vector platforms within the gene therapy landscape. Signal Transduct. Target Ther. 2021, 6, 53, doi:10.1038/s41392-021-00487-6.

7. Wang, J.-H.; Gessler, D.J.; Zhan, W.; Gallagher, T.L.; Gao, G. Adeno-associated virus as a delivery vector for gene therapy of human diseases. Signal Transduct. Target Ther. 2024, 9, 78, doi:10.1038/s41392-024-01780-w.

8. Komatsu, T.; Haruki, H.; Nagata, K. Cellular and viral chromatin proteins are positive factors in the regulation of adenovirus gene expression. Nucleic Acids Res. 2011, 39, 889–901, doi:10.1093/nar/gkq783.

9. Tate, V.E.; Philipson, L. Parental adenovirus DNA accumulates in nucleosome-like structures in infected cells. Nucleic Acids Res. 1979, 6, 2769–2785, doi:10.1093/nar/6.8.2769.

10. Wong, C.M.; McFall, E.R.; Burns, J.K.; Parks, R.J. The role of chromatin in adenoviral vector function. Viruses 2013, 5, 1500–1515, doi:10.3390/v5061500.

11. Herweijer, H.; Wolff, J.A. Progress and prospects: naked DNA gene transfer and therapy. Gene Ther. 2003, 10, 453–458, doi:10.1038/sj.gt.3301983.

12. Togashi, R.; Harashima, H.; Kamiya, H. Correlation between transgen expression and plasmid DNA loss in mouse liver. J. Gene Med. 2013, 15, 242–248, doi:10.1002/jgm.2716.

13. Hobernik, D.; Bros, M. DNA Vaccines—How Far From Clinical Use? J. Mol. Med. 2018, 19, doi:10.3390/ijms19113605.

14. Herweijer, H.; Zhang, G.; Subbotin, V.M.; Budker, V.; Williams, P.; Wolff, J.A. Time course of gene expression after plasmid DNA gene transfer to the liver. J. Gene Med. 2001, 3, 280–291, doi:doi.org/10.1002/jgm.178.

15. Ghazizadeh, S.; Kalish, R.S.; Taichman, L.B. Immune-mediated loss of transgene expression in skin: implications for cutaneous gene therapy. Mol. Ther. 2003, 7, 296–303, doi:10.1016/s1525-0016(03)00013-3.

16. Carroll, A.C.; Wong, A. Plasmid persistence: costs, benefits, and the plasmid paradox. Can. J. Microbiol. 2018, 64, 293–304, doi:10.1139/cjm-2017-0609.

17. Papapetrou, E.P.; Ziros, P.G.; Micheva, I.D.; Zoumbos, N.C.; Athanassiadou, A. Gene transfer into human hematopoietic progenitor cells with an episomal vector carrying an S/MAR element. Gene Ther. 2006, 13, 40–51, doi:10.1038/sj.gt.3302593.

18. Prösch, S.; Stein, J.; Staak, K.; Liebenthal, C.; Volk, H.D.; Krüger, D.H. Inactivation of the very strong HCMV immediate early promoter by DNA CpG methylation in vitro. Biol. Chem. Biol. Chem. Hoppe-Seyler 1996, 377, 195–201, doi:10.1515/bchm3.1996.377.3.195.

19. Hong, K.; Sherley, J.; Lauffenburger, D.A. Methylation of episomal plasmids as a barrier to transient gene expression via a synthetic delivery vector. Biomol. Eng. 2001, 18, 185–192, doi:10.1016/s1389-0344(01)00100-9.

20. Habib, O.; Mohd Sakri, R.; Ghazalli, N.; Chau, D.-M.; Ling, K.-H.; Abdullah, S. Limited expression of non-integrating CpG-free plasmid is associated with increased nucleosome enrichment. PloS One 2020, 15, e0244386, doi:10.1371/journal.pone.0244386.

21. Riu, E.; Chen, Z.-Y.; Xu, H.; He, C.-Y.; Kay, M.A. Histone Modifications are Associated with the Persistence or Silencing of Vector-mediated Transgene Expression <em>In Vivo</em>. Mol. Ther. 2007, 15, 1348–1355, doi:10.1038/sj.mt.6300177.

22. Ochiai, H.; Fujimuro, M.; Yokosawa, H.; Harashima, H.; Kamiya, H. Transient activation of transgene expression by hydrodynamics-based injection may cause rapid decrease in plasmid DNA expression. Gene Ther. 2007, 14, 1152–1159, doi:10.1038/sj.gt.3302970.

23. Kornberg, R.D.; Lorch, Y. Twenty-Five Years of the Nucleosome, Fundamental Particle of the Eukaryote Chromosome. Cell 1999, 98, 285–294, doi:10.1016/s0092-8674(00)81958-3.

24. Liu, R.; Wu, J.; Guo, H.; Yao, W.; Li, S.; Lu, Y.; Jia, Y.; Liang, X.; Tang, J.; Zhang, H. Post-translational modifications of histones: Mechanisms, biological functions, and therapeutic targets. MedComm. 2023, 4, e292, doi:10.1002/mco2.292.

25. Zhang, P.; Torres, K.; Liu, X.; Liu, C.G.; Pollock, R.E. An Overview of Chromatin-Regulating Proteins in Cells. Curr. Protein Pept. Sci. 2016, 17, 401–410, doi:10.2174/1389203717666160122120310.

26. Lee, D.Y.; Hayes, J.J.; Pruss, D.; Wolffe, A.P. A positive role for histone acetylation in transcription factor access to nucleosomal DNA. Cell 1993, 72, 73–84, doi:10.1016/0092-8674(93)90051-q.

27. Kundu, T.K.; Wang, Z.; Roeder, R.G. Human TFIIIC relieves chromatin-mediated repression of RNA polymerase III transcription and contains an intrinsic histone acetyltransferase activity. Mol. Cell Biol. 1999, 19, 1605–1615, doi:10.1128/mcb.19.2.1605.

28. Dion, M.F.; Altschuler, S.J.; Wu, L.F.; Rando, O.J. Genomic characterization reveals a simple histone H4 acetylation code. PNAS 2005, 102, 5501–5506, doi:10.1073/pnas.0500136102.

29. Shogren-Knaak, M.; Ishii, H.; Sun, J.M.; Pazin, M.J.; Davie, J.R.; Peterson, C.L. Histone H4-K16 acetylation controls chromatin structure and protein interactions. Science 2006, 311, 844–847, doi:10.1126/science.1124000.

30. Braunstein, M.; Rose, A.B.; Holmes, S.G.; Allis, C.D.; Broach, J.R. Transcriptional silencing in yeast is associated with reduced nucleosome acetylation. Genes Dev. 1993, 7, 592–604, doi:10.1101/gad.7.4.592.

31. Kouzarides, T. Chromatin Modifications and Their Function. Cell 2007, 128, 693–705, doi:10.1016/j.cell.2007.02.005.

32. Jin, Q.; Yu, L.R.; Wang, L.; Zhang, Z.; Kasper, L.H.; Lee, J.E.; Wang, C.; Brindle, P.K.; Dent, S.Y.R.; Ge, K. Distinct roles of GCN5/PCAF-mediated H3K9ac and CBP/p300-mediated H3K18/27ac in nuclear receptor transactivation. EMBO J. 2011, 30, 249-262-262, doi:doi.org/10.1038/emboj.2010.318.

33. Karmodiya, K.; Krebs, A.R.; Oulad-Abdelghani, M.; Kimura, H.; Tora, L. H3K9 and H3K14 acetylation co-occur at many gene regulatory elements, while H3K14ac marks a subset of inactive inducible promoters in mouse embryonic stem cells. BMC Genom. 2012, 13, 424, doi:10.1186/1471-2164-13-424.

34. Cabrera, A.; Edelstein, H.I.; Glykofrydis, F.; Love, K.S.; Palacios, S.; Tycko, J.; Zhang, M.; Lensch, S.; Shields, C.E.; Livingston, M.; et al. The sound of silence: Transgene silencing in mammalian cell engineering. Cell Syst. 2022, 13, 950–973, doi:10.1016/j.cels.2022.11.005.

35. Zeng, W.; Ball, A.R., Jr.; Yokomori, K. HP1: heterochromatin binding proteins working the genome. Epigenetics 2010, 5, 287–292, doi:10.4161/epi.5.4.11683.

36. Maeda, R.; Tachibana, M. HP1 maintains protein stability of H3K9 methyltransferases and demethylases. EMBO Rep. 2022, 23, e53581, doi:doi.org/10.15252/embr.202153581.

37. Barrett, C.M.; McCracken, R.; Elmer, J.; Haynes, K.A. Components from the Human c-myb Transcriptional Regulation System Reactivate Epigenetically Repressed Transgenes. Int. J. Mol. Sci. 2020, 21, doi:10.3390/ijms21020530.

38. George, O.L.; Ness, S.A. Situational awareness: regulation of the myb transcription factor in differentiation, the cell cycle and oncogenesis. Cancers 2014, 6, 2049–2071, doi:10.3390/cancers6042049.

39. Schmitz, M.L.; Baeuerle, P.A. The p65 subunit is responsible for the strong transcription activating potential of NF-kappa B. Embo J. 1991, 10, 3805–3817, doi:10.1002/j.1460-2075.1991.tb04950.x.

40. Neville, J.J.; Orlando, J.; Mann, K.; McCloskey, B.; Antoniou, M.N. Ubiquitous Chromatin-opening Elements (UCOEs): Applications in biomanufacturing and gene therapy. Biotechnol. Adv. 2017, 35, 557–564, doi:10.1016/j.biotechadv.2017.05.004.

41. Saunders, F.; Sweeney, B.; Antoniou, M.N.; Stephens, P.; Cain, K. Chromatin function modifying elements in an industrial antibody production platform--comparison of UCOE, MAR, STAR and cHS4 elements. PLoS One 2015, 10, e0120096, doi:10.1371/journal.pone.0120096.

42. Sizer, R.E.; White, R.J. Use of ubiquitous chromatin opening elements (UCOE) as tools to maintain transgene expression in biotechnology. Comput. Struct. Biotechnol. J. 2023, 21, 275–283, doi:10.1016/j.csbj.2022.11.059.

43. Betts, Z.; Dickson, A.J. Assessment of UCOE on Recombinant EPO Production and Expression Stability in Amplified Chinese Hamster Ovary Cells. Mol. Biotechnol. 2015, 57, 846–858, doi:10.1007/s12033-015-9877-y.

44. Betts, Z.; Croxford, A.S.; Dickson, A.J. Evaluating the interaction between UCOE and DHFR-linked amplification and stability of recombinant protein expression. Biotechnol. Prog. 2015, 31, 1014–1025, doi:doi.org/10.1002/btpr.2083.

45. Antonova, D.V.; Alekseenko, I.V.; Siniushina, A.K.; Kuzmich, A.I.; Pleshkan, V.V. Searching for Promoters to Drive Stable and Long-Term Transgene Expression in Fibroblasts for Syngeneic Mouse Tumor Models. Int. J. Mol. Sci 2020, 21, doi:10.3390/ijms21176098.

46. Khabou, H.; Cordeau, C.; Pacot, L.; Fisson, S.; Dalkara, D. Dosage Thresholds and Influence of Transgene Cassette in Adeno-Associated Virus-Related Toxicity. Hum. Gene Ther. 2018, 29, 1235–1241, doi:10.1089/hum.2018.144.

47. Sasaki, A.; Kinjo, M. Monitoring intracellular degradation of exogenous DNA using diffusion properties. J. Control. Release 2010, 143, 104–111, doi:doi.org/10.1016/j.jconrel.2009.12.013.

48. Kanazawa, T.; Yamazaki, M.; Fukuda, T.; Takashima, Y.; Okada, H. Versatile nuclear localization signal-based oligopeptide as a gene vector. Biol. Pharm. Bull. 2015, 38, 559–565, doi:10.1248/bpb.b14-00706.

49. Ross, N.L.; Sullivan, M.O. Importin-4 Regulates Gene Delivery by Enhancing Nuclear Retention and Chromatin Deposition by Polyplexes. Mol. Pharm. 2015, 12, 4488–4497, doi:10.1021/acs.molpharmaceut.5b00645.

50. Reilly, M.J.; Larsen, J.D.; Sullivan, M.O. Histone H3 Tail Peptides and Poly(ethylenimine) Have Synergistic Effects for Gene Delivery. Mol. Pharm. 2012, 9, 1031–1040, doi:10.1021/mp200372s.

51. Kamiya, H.; Goto, H.; Kanda, G.; Yamada, Y.; Harashima, H. Transgene expression efficiency from plasmid DNA delivered as a complex with histone H3. Int. J. Pharm. 2010, 392, 249–253, doi:doi.org/10.1016/j.ijpharm.2010.03.035.

52. Kaouass, M.; Beaulieu, R.; Balicki, D. Histonefection: Novel and potent non-viral gene delivery. J. Control. Release 2006, 113, 245–254, doi:doi.org/10.1016/j.jconrel.2006.04.013.

53. Uetsuki, T.; Naito, A.; Nagata, S.; Kaziro, Y. Isolation and Characterization of the Human Chromosomal Gene for Polypeptide Chain Elongation Factor-1α. J. Biol. Chem. 1989, 264, 5791–5798, doi:doi.org/10.1016/S0021-9258(18)83619-5.

54. Wakabayashi-Ito, N.; Nagata, S. Characterization of the regulatory elements in the promoter of the human elongation factor-1 alpha gene. J. Biol. Chem. 1994, 269, 29831–29837, doi:doi.org/10.1016/S0021-9258(18)43956-7.

55. Haase, R.; Argyros, O.; Wong, S.-P.; Harbottle, R.P.; Lipps, H.J.; Ogris, M.; Magnusson, T.; Pinto, M.G.V.; Haas, J.; Baiker, A. pEPito: a significantly improved non-viral episomal expression vector for mammalian cells. BMC Biotechnol. 2010, 10, 20, doi:10.1186/1472-6750-10-20.

56. Kreppel, F.; Hagedorn, C. Episomes and Transposases—Utilities to Maintain Transgene Expression from Nonviral Vectors. Genes 2022, 13, doi:10.3390/genes13101872.

57. Mehta, A.K.; Majumdar, S.S.; Alam, P.; Gulati, N.; Brahmachari, V. Epigenetic regulation of cytomegalovirus major immediate-early promoter activity in transgenic mice. Gene 2009, 428, 20–24, doi:10.1016/j.gene.2008.09.033.

58. Lowary, P.T.; Widom, J. New DNA sequence rules for high affinity binding to histone octamer and sequence-directed nucleosome positioning. J. Mol. Biol. 1998, 276, 19–42, doi:10.1006/jmbi.1997.1494.

59. Jimenez-Useche, I.; Yuan, C. The Effect of DNA CpG Methylation on the Dynamic Conformation of a Nucleosome. Biophys. J. 2012, 103, 2502–2512, doi:doi.org/10.1016/j.bpj.2012.11.012.

60. Nurse, N.P.; Jimenez-Useche, I.; Smith, I.T.; Yuan, C. Clipping of Flexible Tails of Histones H3 and H4 Affects the Structure and Dynamics of the Nucleosome. Biophys. J. 2013, 104, 1081–1088, doi:doi.org/10.1016/j.bpj.2013.01.019.

61. Nurse, N.P.; Yuan, C. Cis and trans internucleosomal interactions of H3 and H4 tails in tetranucleosomes. Biopolymers 2015, 103, 33–40, doi:doi.org/10.1002/bip.22560.

62. Bednar, J.; Horowitz, R.A.; Grigoryev, S.A.; Carruthers, L.M.; Hansen, J.C.; Koster, A.J.; Woodcock, C.L. Nucleosomes, linker DNA, and linker histone form a unique structural motif that directs the higher-order folding and compaction of chromatin. PNAS 1998, 95, 14173–14178, doi:10.1073/pnas.95.24.14173.

63. Cleri, F.; Giordano, S.; Blossey, R. Nucleosome Array Deformation in Chromatin is Sustained by Bending, Twisting and Kinking of Linker DNA. J Mol Biol. 2023, 435, 168263, doi:doi.org/10.1016/j.jmb.2023.168263.

64. Jimenez-Useche, I.; Nurse, N.P.; Tian, Y.; Kansara, B.S.; Shim, D.; Yuan, C. DNA methylation effects on tetra-nucleosome compaction and aggregation. Biophys. J. 2014, 107, 1629–1636, doi:10.1016/j.bpj.2014.05.055.

65. Carroll, D. Continuous-flow salt gradient dialysis for the preparation of polynucleotide-polypeptide complexes. Anal. Biochem. 1971, 44, 496–502, doi:doi.org/10.1016/0003-2697(71)90237-5.

66. Jordan, M.; Schallhorn, A.; Wurm, F.M. Transfecting mammalian cells: Optimization of critical parameters affecting calcium-phosphate precipitate formation. Nucleic Acids Res. 1996, 24, 596–601, doi:10.1093/nar/24.4.596.

67. Bewick, V.; Cheek, L.; Ball, J. Statistics review 9: One-way analysis of variance. Crit. Care 2004, 8, 130, doi:10.1186/cc2836.

68. Abdi, H.; Williams, L.J. Tukey’s honestly significant difference (HSD) test. Encyclopedia of Research Design 2010, 3, 1–5.

69. Valouev, A.; Johnson, S.M.; Boyd, S.D.; Smith, C.L.; Fire, A.Z.; Sidow, A. Determinants of nucleosome organization in primary human cells. Nature 2011, 474, 516–520, doi:10.1038/nature10002.

70. Makde, R.D.; England, J.R.; Yennawar, H.P.; Tan, S. Structure of RCC1 chromatin factor bound to the nucleosome core particle. Nature 2010, 467, 562–566, doi:10.1038/nature09321.

71. Zhao, H.; Guo, M.; Zhang, F.; Shao, X.; Liu, G.; Xing, Y.; Zhao, X.; Luo, L.; Cai, L. Nucleosome Assembly and Disassembly in vitro Are Governed by Chemical Kinetic Principles. Front. Cell. Dev. Biol. 2021, 9, 762571, doi:10.3389/fcell.2021.762571.

72. Flaus, A. Principles and practice of nucleosome positioning in vitro. Front. Life Sci. 2011, 5, 5–27, doi:10.1080/21553769.2012.702667.

73. Wang, X.; Xu, Z.; Tian, Z.; Zhang, X.; Xu, D.; Li, Q.; Zhang, J.; Wang, T. The EF-1α promoter maintains high-level transgene expression from episomal vectors in transfected CHO-K1 cells. J. Cell. Mol. Med. 2017, 21, 3044–3054, doi:10.1111/jcmm.13216.

74. Rogge, R.A.; Kalashnikova, A.A.; Muthurajan, U.M.; Porter-Goff, M.E.; Luger, K.; Hansen, J.C. Assembly of nucleosomal arrays from recombinant core histones and nucleosome positioning DNA. J. Vis. Exp. 2013, doi:10.3791/50354.

75. Vaquero, A.; Scher, M.B.; Lee, D.H.; Sutton, A.; Cheng, H.L.; Alt, F.W.; Serrano, L.; Sternglanz, R.; Reinberg, D. SirT2 is a histone deacetylase with preference for histone H4 Lys 16 during mitosis. Genes Dev. 2006, 20, 1256–1261, doi:10.1101/gad.1412706.

76. Chen, H.P.; Zhao, Y.T.; Zhao, T.C. Histone deacetylases and mechanisms of regulation of gene expression. Crit. Rev. Oncog. 2015, 20, 35–47, doi:10.1615/critrevoncog.2015012997.

77. Shabane, P.S.; Onufriev, A.V. Significant compaction of H4 histone tail upon charge neutralization by acetylation and its mimics, possible effects on chromatin structure. J. Mol. Biol. 2021, 433, 166683, doi:10.1016/j.jmb.2020.10.017.

78. Wang, X.; Hayes, J.J. Acetylation mimics within individual core histone tail domains indicate distinct roles in regulating the stability of higher-order chromatin structure. Mol. Cell. Biol. 2008, 28, 227–236, doi:10.1128/mcb.01245-07.

79. Mishima, Y.; Miyagi, S.; Saraya, A.; Negishi, M.; Endoh, M.; Endo, T.A.; Toyoda, T.; Shinga, J.; Katsumoto, T.; Chiba, T.; et al. The Hbo1-Brd1/Brpf2 complex is responsible for global acetylation of H3K14 and required for fetal liver erythropoiesis. Blood 2011, 118, 2443–2453, doi:10.1182/blood-2011-01-331892.

80. Wang, Z.; Zang, C.; Rosenfeld, J.A.; Schones, D.E.; Barski, A.; Cuddapah, S.; Cui, K.; Roh, T.Y.; Peng, W.; Zhang, M.Q.; et al. Combinatorial patterns of histone acetylations and methylations in the human genome. Nat. Genet. 2008, 40, 897–903, doi:10.1038/ng.154.

81. Christensen, M.D.; Nitiyanandan, R.; Meraji, S.; Daer, R.; Godeshala, S.; Goklany, S.; Haynes, K.; Rege, K. An inhibitor screen identifies histone-modifying enzymes as mediators of polymer-mediated transgene expression from plasmid DNA. J. Control Release 2018, 286, 210–223, doi:10.1016/j.jconrel.2018.06.030.

82. Elmer, J.J.; Christensen, M.D.; Barua, S.; Lehrman, J.; Haynes, K.A.; Rege, K. The histone deacetylase inhibitor Entinostat enhances polymer-mediated transgene expression in cancer cell lines. Biotechnol. Bioeng. 2016, 113, 1345–1356, doi:10.1002/bit.25898.

83. Barua, S.; Rege, K. The influence of mediators of intracellular trafficking on transgene expression efficacy of polymer-plasmid DNA complexes. Biomaterials 2010, 31, 5894–5902, doi:10.1016/j.biomaterials.2010.04.007.

84. Struhl, K.; Segal, E. Determinants of nucleosome positioning. Nat. Struct. Mol. Biol. 2013, 20, 267–273, doi:10.1038/nsmb.2506.

85. Gu, S.G.; Fire, A. Partitioning the C. elegans genome by nucleosome modification, occupancy, and positioning. Chromosoma 2010, 119, 73–87, doi:10.1007/s00412-009-0235-3.

86. Saxton, D.S.; Rine, J. Nucleosome Positioning Regulates the Establishment, Stability, and Inheritance of Heterochromatin in Saccharomyces cerevisiae. PNAS 2020, 117, 27493–27501, doi:10.1073/pnas.2004111117.

87. Taylor, G.C.; Eskeland, R.; Hekimoglu-Balkan, B.; Pradeepa, M.M.; Bickmore, W.A. H4K16 acetylation marks active genes and enhancers of embryonic stem cells, but does not alter chromatin compaction. Genome Res. 2013, 23, 2053–2065, doi:10.1101/gr.155028.113.

88. Reeves, R.; Gorman, C.M.; Howard, B. Minichromosome assembly of non-integrated plasmid DNA transfected into mammalian cells. Nucleic Acids Res. 1985, 13, 3599–3615, doi:10.1093/nar/13.10.3599.

89. O’Connor, L.; Gilmour, J.; Bonifer, C. The Role of the Ubiquitously Expressed Transcription Factor Sp1 in Tissue-specific Transcriptional Regulation and in Disease. Yale J. Biol. Med. 2016, 89, 513–525.

90. Wu, Z.; Nicoll, M.; Ingham, R.J. AP-1 family transcription factors: a diverse family of proteins that regulate varied cellular activities in classical hodgkin lymphoma and ALK+ ALCL. Exp. Hematol. Oncol. 2021, 10, 4, doi:10.1186/s40164-020-00197-9.

91. Elgin, S.C.; Reuter, G. Position-effect variegation, heterochromatin formation, and gene silencing in Drosophila. Cold Spring Harb. Perspect. Biol. 2013, 5, a017780, doi:10.1101/cshperspect.a017780.

92. Gates, L.A.; Shi, J.; Rohira, A.D.; Feng, Q.; Zhu, B.; Bedford, M.T.; Sagum, C.A.; Jung, S.Y.; Qin, J.; Tsai, M.J.; et al. Acetylation on histone H3 lysine 9 mediates a switch from transcription initiation to elongation. J. Biol. Chem. 2017, 292, 14456–14472, doi:10.1074/jbc.M117.802074.

93. Strober, W. Trypan Blue Exclusion Test of Cell Viability. Curr. Protoc. Immunol. 2015, 111, A3.b.1-a3.b.3, doi:10.1002/0471142735.ima03bs111.

94. Li, C.; Goryaynov, A.; Yang, W. The selective permeability barrier in the nuclear pore complex. Nucleus 2016, 7, 430–446, doi:10.1080/19491034.2016.1238997.

95. van der Aa, M.A.; Mastrobattista, E.; Oosting, R.S.; Hennink, W.E.; Koning, G.A.; Crommelin, D.J. The nuclear pore complex: the gateway to successful nonviral gene delivery. Pharm Res. 2006, 23, 447–459, doi:10.1007/s11095-005-9445-4.

96. Shimozono, S.; Tsutsui, H.; Miyawaki, A. Diffusion of large molecules into assembling nuclei revealed using an optical highlighting technique. Biophys J. 2009, 97, 1288–1294, doi:10.1016/j.bpj.2009.06.024.

97. Bernardes, N.E.; Fung, H.Y.J.; Li, Y.; Chen, Z.; Chook, Y.M. Structure of IMPORTIN-4 bound to the H3–H4– ASF1 histone–histone chaperone complex. PNAS 2022, 119, e2207177119, doi:10.1073/pnas.2207177119.

98. Sumida, N.; Nishikawa, J.; Kishi, H.; Amano, M.; Furuya, T.; Sonobe, H.; Ohyama, T. A designed curved DNA segment that is a remarkable activator of eukaryotic transcription. FEBS J. 2006, 273, 5691–5702, doi:10.1111/j.1742-4658.2006.05557.x.

99. Kamiya, H.; Fukunaga, S.; Ohyama, T.; Harashima, H. The location of the left-handedly curved DNA sequence affects exogenous DNA expression in vivo. Arch. Biochem. Biophys. 2007, 461, 7–12, doi:10.1016/j.abb.2007.02.012.

100. Kamiya, H.; Goto, H.; Harashima, H. Effects of non-B DNA sequences on transgene expression. J Biosci. Bioeng. 2009, 108, 20–23, doi:10.1016/j.jbiosc.2009.02.013.

101. Zhang, J.; Cheng, S.; Yang, W.; Li, S. Enhanced transgene expression using two β-globin MARs flanking expression cassettes in stably transfected CHO-K1 cells. 3 Biotech. 2019, 9, 435, doi:10.1007/s13205-019-1971-6.

## References

1. Lowary, P.T.; Widom, J. New DNA sequence rules for high affinity binding to histone octamer and sequence-directed nucleosome positioning. J. Mol. Biol. 1998, 276, 19–42, doi:10.1006/jmbi.1997.1494.

2. Makde, R.D.; England, J.R.; Yennawar, H.P.; Tan, S. Structure of RCC1 chromatin factor bound to the nucleosome core particle. Nature 2010, 467, 562–566, doi:10.1038/nature09321.

3. Uetsuki, T.; Naito, A.; Nagata, S.; Kaziro, Y. Isolation and Characterization of the Human Chromosomal Gene for Polypeptide Chain Elongation Factor-1α. J. Biol. Chem. 1989, 264, 5791–5798, doi:doi.org/10.1016/S0021-9258(18)83619-5.

4. Wakabayashi-Ito, N.; Nagata, S. Characterization of the regulatory elements in the promoter of the human elongation factor-1 alpha gene. J. Biol. Chem. 1994, 269, 29831–29837, doi:doi.org/10.1016/S0021-9258(18)43956-7.

5. Jimenez-Useche, I.; Yuan, C. The Effect of DNA CpG Methylation on the Dynamic Conformation of a Nucleosome. Biophys. J. 2012, 103, 2502–2512, doi:doi.org/10.1016/j.bpj.2012.11.012.

6. Nurse, N.P.; Yuan, C. Cis and trans internucleosomal interactions of H3 and H4 tails in tetranucleosomes. Biopolymers 2015, 103, 33–40, doi:doi.org/10.1002/bip.22560.

7. Bewick, V.; Cheek, L.; Ball, J. Statistics review 9: One-way analysis of variance. Crit. Care 2004, 8, 130, doi:10.1186/cc2836.

8. Abdi, H.; Williams, L.J. Tukey’s honestly significant difference (HSD) test. Encyclopedia of Research Design 2010, 3, 1–5.

